# Dual Activation of MC3R and MC4R Drives Weight Loss and Reduces Food Intake in Obese Primates

**DOI:** 10.1101/2025.11.25.690554

**Authors:** Jillian L Seiler, Anna C Impastato, Emma (Xiaoyu) Zhang, Kade J Kelley, Thomas L Bennett, Bradley Studnitzer, Claudia R Prindle, Benjamin H Rajewski, Barry A Badeau, Xinjian Jiang, Russell Potterfield, Jordan Y Delev, Daniel L Marks

## Abstract

The melanocortin system plays a central role in regulating hunger and satiety, making it an attractive target for treating metabolic disease. However, the limited clinical success of selective melanocortin-4 receptor (MC4R) agonists prompted the investigation of whether concurrent melanocortin-3 receptor (MC3R) and MC4R activation is key to unlocking the melanocortin system for the treatment of general obesity. To test this hypothesis, we designed and synthesized novel peptides to probe the distinct and combined roles of MC3R and MC4R in nonhuman primates (NHPs). We show that selectively agonizing MC3R modulates food intake in a state-dependent manner. Moreover, co-agonism of MC3R and MC4R results in more substantial metabolic effects than selective MC4R agonism, highlighting both a non-redundant and a cooperative role of MC3R. To leverage these discoveries, we developed 710GO, an orally-available MC3R/MC4R dual agonist peptide that induces significant weight loss in diet-induced obese (DIO) NHPs. Oral 710GO treatment demonstrates limited weight rebound, has additive effects in combination with GLP-1s, and exhibits a clean safety profile. These results reestablish the melanocortin system, specifically concerted MC3R/MC4R agonism, as a viable mechanism for next-generation obesity therapeutics.

## Main

The recent successes of GLP-1s in treating obesity address a previously unmet demand for weight loss therapeutics. However, side effects and variable efficacy underscore the need for mechanistically-differentiated approaches to support a massive and diverse patient population.^1–3^ The melanocortin (MC) family of receptors have long represented an alternative metabolic target to GLP-1s for treating obesity, but drugs targeting MC receptors are currently limited to treating obesity caused by rare genetic conditions.^4,5^ A greater understanding of the MC family is required to unlock its potential for weight loss in general obesity populations.

The MC family of receptors is a highly conserved neuroendocrine signaling network that plays a critical role in energy homeostasis.^6–10^ It is comprised of five G protein-coupled receptors (MC1R-MC5R) that are endogenously activated by peptides derived from the proopiomelanocortin (POMC) precursor polypeptide—α-, β-, γ-MSH and ACTH.^11^ One well-known endogenous antagonist of the MC family, Agouti-related peptide (AgRP), acts primarily on two receptors—MC3R and MC4R.^12^ These are the most highly expressed MCRs in the brain, with significant expression in areas of the hypothalamus that regulate food intake and peripheral metabolism (e.g., the arcuate nucleus (ARC) and paraventricular nucleus (PVN)).^6,13^ MC3R is expressed presynaptically on POMC/AgRP terminals that regulate GABA-mediated inhibition onto postsynaptic MC4R-expressing PVN neurons, highlighting the direct ability for MC3R to modulate MC4R activity.^14^ While agonism of the central melanocortin system (e.g., α-MSH) causes reduced food intake and weight loss, antagonism (e.g. AgRP) causes increased food intake and weight gain.^15^ Notably, both of these peptides simultaneously target MC3R and MC4R. Assuming these two receptors have non-redundant functions, their shared ligands and hypothalamic expression pattern indicate a potentially cooperative role in metabolism.

Despite evidence for these receptors’ synergy, early rodent studies predominately suggest the paramount role of MC4R over MC3R in body weight regulation.^16–20^ Furthermore, MC4R loss-of-function (LoF) mutations were identified as the most common cause of monogenic obesity in humans.^21,22^ These observations, combined with the limited pharmacological evaluation of MC3R, likely primed pharmaceutical efforts to focus on MC4R-selective agonists for treating obesity. All of these MC4R-selective programs were terminated after failing to demonstrate safe, translational efficacy in the clinic. We hypothesize these compounds had insufficient therapeutic windows, wherein adverse effects prevented dosing at efficacious levels in humans.

Recently, mifomelatide (TCMCB07), a first-in-class MC3R/MC4R dual antagonist we developed to treat involuntary weight loss, showed increases in participant body weight in a phase I clinical trial.^23–26^ Due to its translational efficacy and its dual antagonism on MC3R and MC4R, we decided to investigate MC3R/MC4R dual agonism as a potential mechanism for safe, efficacious weight loss in humans.

To evaluate the effects of MC3R and MC4R on food intake and body weight, we designed and synthesized a series of peptides with differing selectivity profiles. The compounds were then evaluated in functional assays measuring cyclic adenosine monophosphate (cAMP) production (**Fig. 1**, **Table 1, Supplementary Fig. 1**). Functional activity of α-MSH was comparable to reported values, validating the assay.^27^ All previous clinical molecules demonstrated full MC4R agonism (E_max_ ≥ 100%) and selectivity for MC4R over MC3R. Additionally, we designed MC3R- and MC4R-selective compounds (EBMC-40-187 and EBMC-03-47, respectively) and compounds with full agonist activity on both MC3R and MC4R (710GO) and mixed activity as an MC3R partial antagonist/MC4R full agonist (707GO). We additionally measured the binding affinity on MC3R and MC4R and calculated IC_50_ values for 710GO and 707GO (**Supplementary Fig. 2)**.

**Fig. 1.**
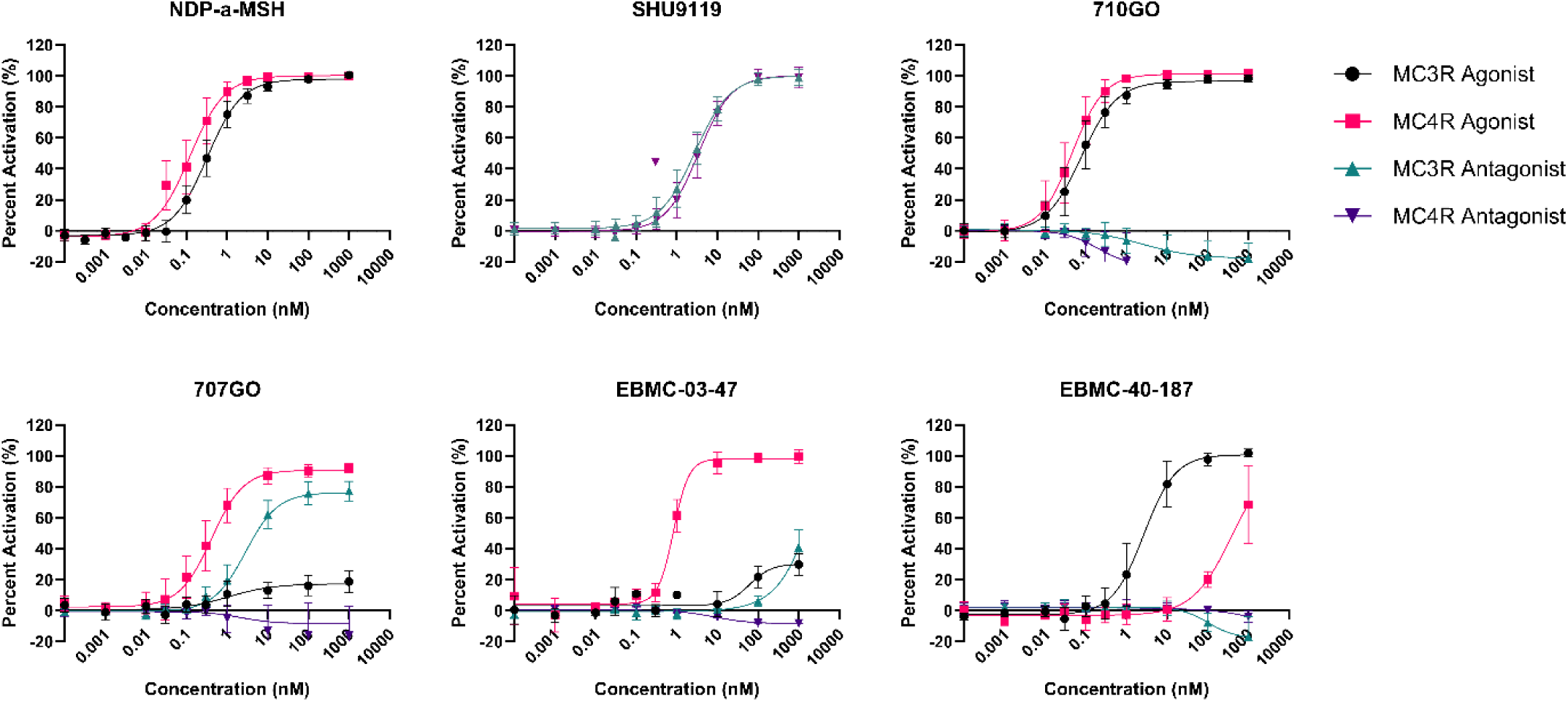
Pharmacological characterization of melanocortin ligands against human MC3R and human MC4R expressed in CHO-K1 cells by cAMP signaling. Percent activation is relative to NDP-a-MSH for agonist mode and SHU 9119 for antagonist mode.

**Table 1.** Melanocortin ligands and their corresponding agonist and antagonist potencies on MC3R and MC4R. (EC_50_, effective concentration at 50% activation; E_max_, response relative to NDP-α-MSH; n.t., not tested)

| Company | Compound ID | cAMP Signaling<br>EC<sub>50</sub> (nM) Emax (%) |  |  |  |
| --- | --- | --- | --- | --- | --- |
|  |  | Agonist |  | Antagonist |  |
|  |  | MC3R | MC4R | MC3R | MC4R |
| **Endogenous Ligands** | α-MSH | 0.57 (102%) | 0.49 (99%) | >1,000 | >1,000 |
| | β-MSH | 0.31 (96%) | 0.35 (100%) | >1,000 | >1,000 |
| | γ-1-MSH | 0.07 (101%) | 1.01 (96%) | >1,000 | >1,000 |
|  | AgRP | >1,000 | >1,000 | 3.19 (103%) | 31.0 (97%) |
| **Endevica Bio** | mifomelatide | >1,000 | >1,000 | 3.25 (95%) | 7.24 (98%) |
|  | 707GO | 250 (25%) | 0.36 (90%) | 2.39 (77%) | >1,000 |
|  | 710GO | 0.073 (96%) | 0.042 (101%) | >1,000 | >1,000 |
|  | EBMC-03-47 | >1,000 | 0.89 (99%) | >1,000 | >1,000 |
|  | EBMC-40-187 | 2.6 (100%) | 339 (100%) | >1,000 | >1,000 |
| **Palatin** | bremelanotide | 5.5 (92%) | 0.22 (99%) | >1,000 | >1,000 |
| **Clinuvel** | NDP-α-MSH | 0.32 (98%) | 0.17 (100%) | n.t. | n.t. |
| **Novo Nordisk** | MC4-NN2-0453 | 39 (31%) | 31 (105%) | >1,000 | >1,000 |
| **Eli Lilly** | LY2112688 | 4.2 (91%) | 0.048 (100%) | >1,000 | >1,000 |
| **Merck** | MK-0493 | >1,000 | 7.8 (104%) | >1,000 | >1,000 |
| **Rhythm** | bivamelagon | 130 (102%) | 0.98 (103%) | >1,000 | >1,000 |
|  | setmelanotide | 0.82 (88%) | 0.053 (100%) | >1,000 | >1,000 |

MC3R is implicated as a key regulator of energy homeostasis and is critical for providing an upper bound to weight gain during exposure to energy-dense food.^14^ To investigate the roles of MC3R and MC4R in the context of homeostatic bounds, we subcutaneously administered MC4R-selective agonist EBMC-03-47, MC3R-selective agonist EBMC-40-187, and their combination to male NHPs in different metabolic states: *ad libitum*/lean and fasted/lean. Food intake was measured over 24 hours following drug administration and was normalized to their own vehicle baseline.

In lean animals that were fed *ad libitum*, MC3R or MC4R agonism did not significantly alter food intake compared to control after 24 hours (**Fig. 2A**). Total food consumption during the first two hours was highly variable but low, leading to large changes in cumulative food intake that normalized after three hours (**Supplementary Fig. 3**). In NHPs that were fasted overnight, treatment with selective agonists alone and in combination resulted in short-term decreases in food intake (**Fig. 2B**). MC4R agonism reduced food intake by 20.0 ± 2.4% in the first hour after dosing but returned to baseline after four hours. The effect of MC3R agonism was more pronounced, reducing cumulative food intake by 46.0 ± 7.1% in the first hour. This reduction was persistent; after 24 hours, cumulative food intake was still 14.3 ± 1.6% lower than control. Co-administration of both agonists caused the largest reduction of cumulative food intake for the first three hours, averaging 54.9 ± 18.9% lower than control. However, these levels returned to baseline after four hours. Compared to *ad libitum* conditions, modest homeostatic challenge (fasting) increased the food intake reduction of melanocortin agonism. Further, the enduring cumulative food intake reduction observed with MC3R agonist treatment, in contrast to MC4R-selective and combination treatment, supports the claim that MC3R provides important boundary control in metabolic processes.

**Fig. 2.**
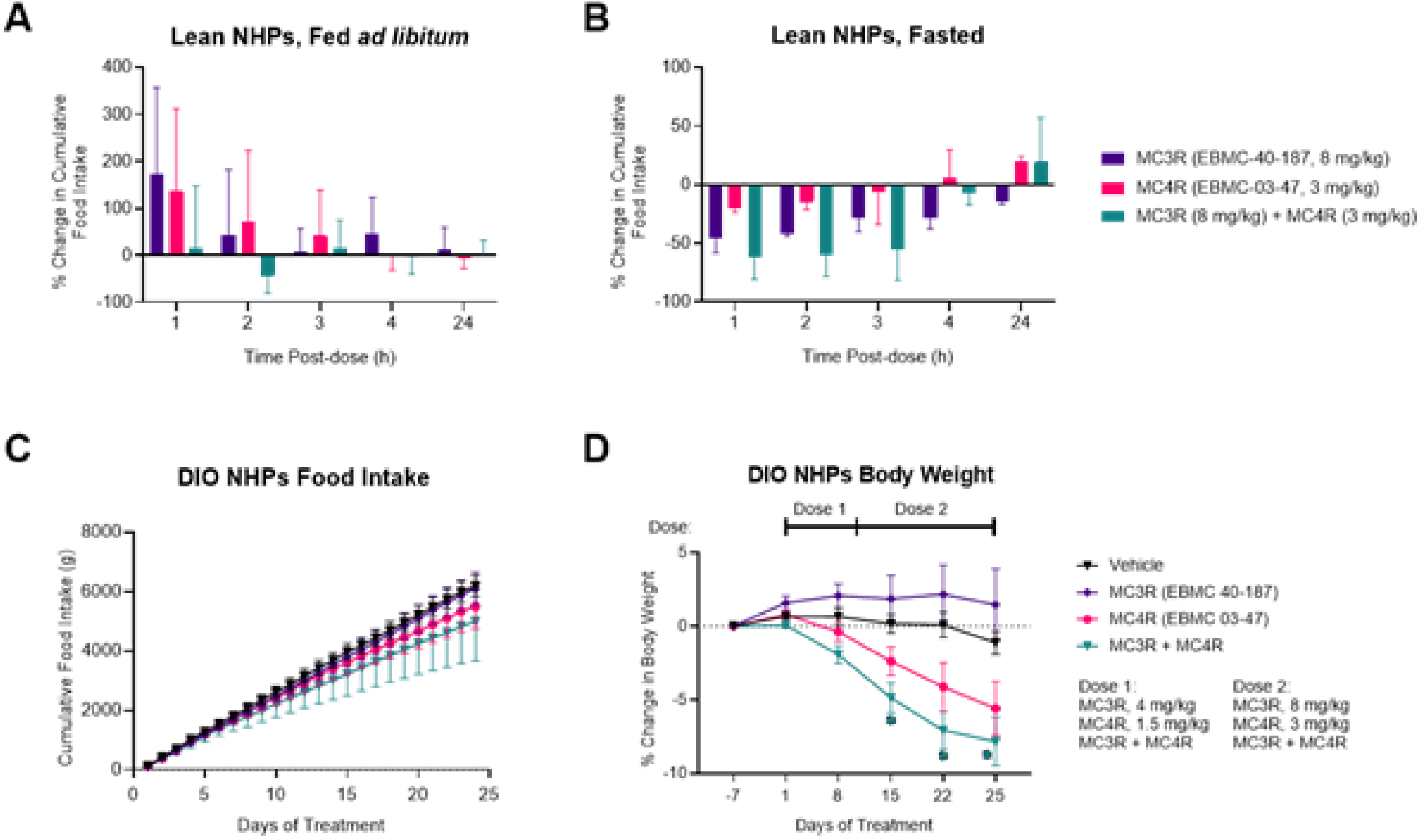
Effects of MC3R, MC4R, and MC3R + MC4R co-agonism on food intake and body weight in male NHPs. **A,B)** Change in cumulative food intake in *ad libitum*, lean male NHPs (A) and fasted, lean male NHPs (B) normalized to placebo (n=3, per group). **C,D)** Individual and combined effects of MC3R and MC4R agonism on cumulative food intake (C) and percent change in body weight (D) in DIO male NHPs, normalized to each individual initial body weight. Animals were administered vehicle (300 mM mannitol in sWFI, inverted black triangle) or test compound by subcutaneous injection (n=5, per dose group; n=4 for co-agonism group). EBMC-40-187 (purple diamond) is a full MC3R agonist, EBMC-03-47 (pink circle) is a full MC4R agonist, and the two peptides were co-administered for MC3R + MC4R co-agonism (inverted green triangle). In the acute lean NHP study, EBMC-03-47 was administered at 3 mg/kg, SC and EBMC-40-187 was administered at 8 mg/kg, SC (A, B). In the subchronic DIO-NHP study, the EBMC-03-47 dose was increased from 1.5 mg/kg to 3 mg/kg and the EBMC-40-187 dose was increased from 4 mg/kg to 8 mg/kg on Day 11 in both the individual and combination treatment groups (C,D). All data are presented as mean ± SEM. Statistical significance was tested where appropriate using Two-Way ANOVA with Dunnett’s multiple comparison test. *p < 0.05

We then investigated the effect of repeated dosing of MC3R/MC4R agonists in DIO NHPs. An MC3R agonist, MC4R agonist, and a combination of the two were administered subcutaneously to DIO NHPs daily. These animals were provided with *ad libitum* high-fat diet, and food intake and body weight were measured daily for 15 days. On day ten, doses were increased to investigate dose responsiveness. Over the duration of the study, cumulative food intake decreased across all groups compared to vehicle treatment, although with large individual variability (**Fig. 2C**). MC3R agonism decreased cumulative food intake by 1.4 ± 8.9% compared to vehicle, whereas MC4R agonism reduced food intake by 11.2 ± 12.1%. Consistent with the lean fasted NHPs in the acute feeding study, co-administration of MC3R and MC4R agonists resulted in the greatest reduction in cumulative food intake—a 19.6 ± 21.3% decrease compared to vehicle. These results point to a cooperative effect of MC3R and MC4R agonism on food intake in DIO NHPs consuming an energy-dense diet.

The observed food intake reduction from MC3R/MC4R co-agonism was well coupled with decreases in body weight (**Fig. 2D, Supplementary Fig. 3**). Co-administration of selective agonists significantly reduced body weight by 7.8 ± 1.7% over the course of the study (p<0.05). This decrease in body weight was substantial compared to selective MC4R agonism, which only reduced body weight by 5.6 ± 1.8%. Interestingly, selective MC3R agonism resulted in 1.5 ± 2.4% *increase* in body weight, despite slightly reducing food intake. While MC4R agonism is sufficient for weight loss, these data suggest that co-agonism with MC3R leads to greater weight loss. Comparable results were obtained in rats, with the notable exception that selective MC3R agonism elicited weight loss (**Supplementary Figs. 4 & 7**). This species-specific divergence may reflect evolutionary differences in melanocortin-mediated metabolic regulation.

While our data suggests complementary MC3R/MC4R co-agonism will improve weight loss, other literature demonstrated that *de-activating* MC3R (through pharmacological antagonism or gene knockout) can facilitate weight loss and inhibit weight regain in lean rodents.^14,28,29^ These contradictory proposed roles of MC3R necessitated further investigation of MC3R activity in conjunction with MC4R agonism by a single peptide. To evaluate this, we subcutaneously administered two compounds—710GO and 707GO—to fasted DIO NHPs. While both compounds are full MC4R agonists with similar pharmacokinetic (PK) properties, 710GO is a potent MC3R agonist while 707GO is a partial MC3R antagonist (**Table 1, Supplementary Tables 2-4, Supplementary Figs. 9 and 10**). Food intake was measured for 72 hours after subcutaneous injection and plotted alongside the animals’ respective vehicle baseline (**Fig. 3A**). NHPs treated with 710GO experienced a non-significant reduction in food intake that peaked at 0.5 hours post-dose but was still 42.5 ± 8.0% less than vehicle by 72 hours (**Fig. 3B**). NHPs administered 707GO briefly trended towards an increase in food intake but normalized back to vehicle levels of cumulative food intake by 72 hours. We hypothesize that 707GO did not reduce food intake due to either its lack of MC3R agonism or a counteractive effect by its less-potent partial antagonism of MC3R.

**Fig. 3.**
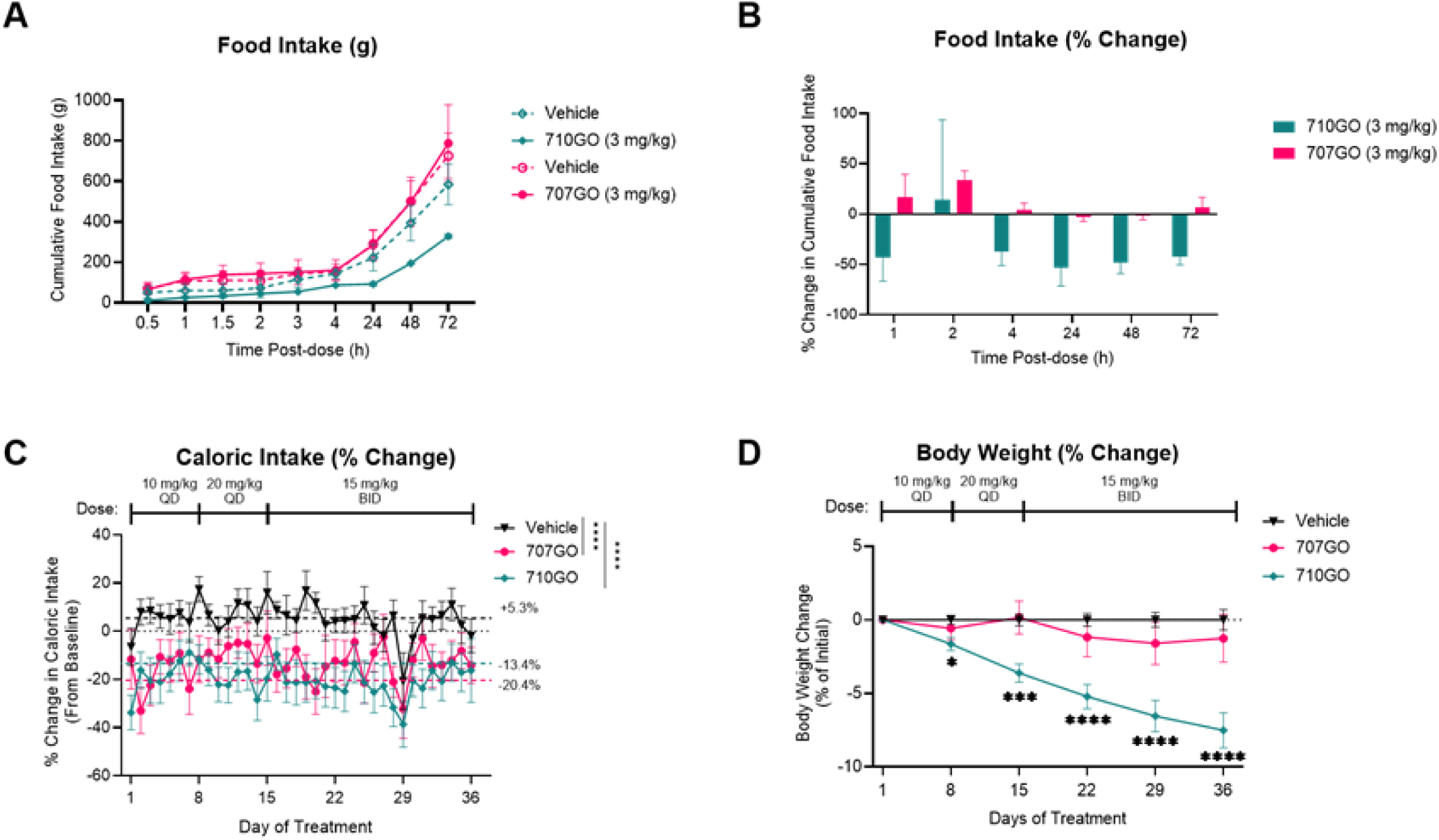
MC3R agonism alters the metabolic effects of MC4R agonism in DIO NHPs. **A)** Cumulative food intake of fasted DIO NHPs (n=2) after administration of vehicle (empty green diamond and empty pink circle, saline) and test compounds 707GO (pink circle, a full MC4R agonist/MC3R partial antagonist) and 710GO (green diamond, a full MC3/4R agonist). **B)** Percent change in cumulative food intake between test compound and vehicle (black inverted triangle) in fasted DIO NHPs (n=2). **C)** Percent change in daily caloric intake and **D)** body weight in DIO NHPs (n=8) during repeated dosing. A single 3 mg/kg SC injection was administered in the acute studies. In the subchronic study, compounds were administered via oral gavage in a scheduled dose-escalation routine: *Days 1-8*: 10 mg/kg, QD; *Days 9-14*: 20 mg/kg, QD; *Days 15-36:* 15 mg/kg, BID. All data are presented as mean ± SEM. Statistical significance in (C) using mixed-effects analysis and (D) using Two-Way ANOVA, both followed by Dunnett’s multiple comparison test (*p < 0.05, ***p<0.001, ****p<0.0001).

We then set out to determine whether this acute interaction between MC3R and MC4R translates to body weight changes with repeated dosing. We administered 710GO and 707GO daily via oral gavage to DIO NHPs for 35 days. NHPs treated with 710GO consumed 20.3 ± 1.1% fewer calories compared to their pre-treatment baseline, whereas NHPs treated with 707GO consumed 13.4 ± 1.3% fewer calories (**Fig. 3C**). Both treatment groups showed statistically significant reductions in caloric consumption across the study duration (p<0.0001), as well as on specific days—with 710GO treated animals showing more days of significant difference. Consistent with monoagonist experiments, subchronic studies in rats show a reduction in food intake and body weight in response to both 710GO and 707GO (**Supplementary Table 1, Supplementary Figs. 5, 7-8**). In NHPs, by day 36, 710GO-treated animals had lost 7.5 ± 1.2% body weight compared to vehicle, but 707GO-treated animals did not demonstrate significant weight change (**Fig. 3D, Supplementary Fig. 6B**). These results further suggest that MC3R agonism works cooperatively with MC4R agonism to produce larger reductions in body weight than selective MC4R agonism.

After establishing the efficacy of dual MC3R/MC4R agonism for weight loss in obese NHPs, we investigated the effects of chronic administration. DIO NHPs were given 710GO (10 mg/kg) or saline by oral gavage. Food intake and body weight measurements were taken daily. DIO NHPs were offered three meals per day, including high-fat chow, fruit, and normal chow. Due to the high-calorie diet provided, DIO NHPs receiving saline gained 4.6 ± 0.4% body weight over the course of the study. Interestingly, we noted that 710GO-treated animals shifted preference from highly palatable foods towards the normal chow diet (data not shown). 710GO induced significant weight loss by day 21 and ultimately produced 11.8 ± 2.1% weight loss in 15 weeks compared to vehicle (p<0.05; **Fig. 4A, Supplementary Figure 6C**).

**Fig. 4.**
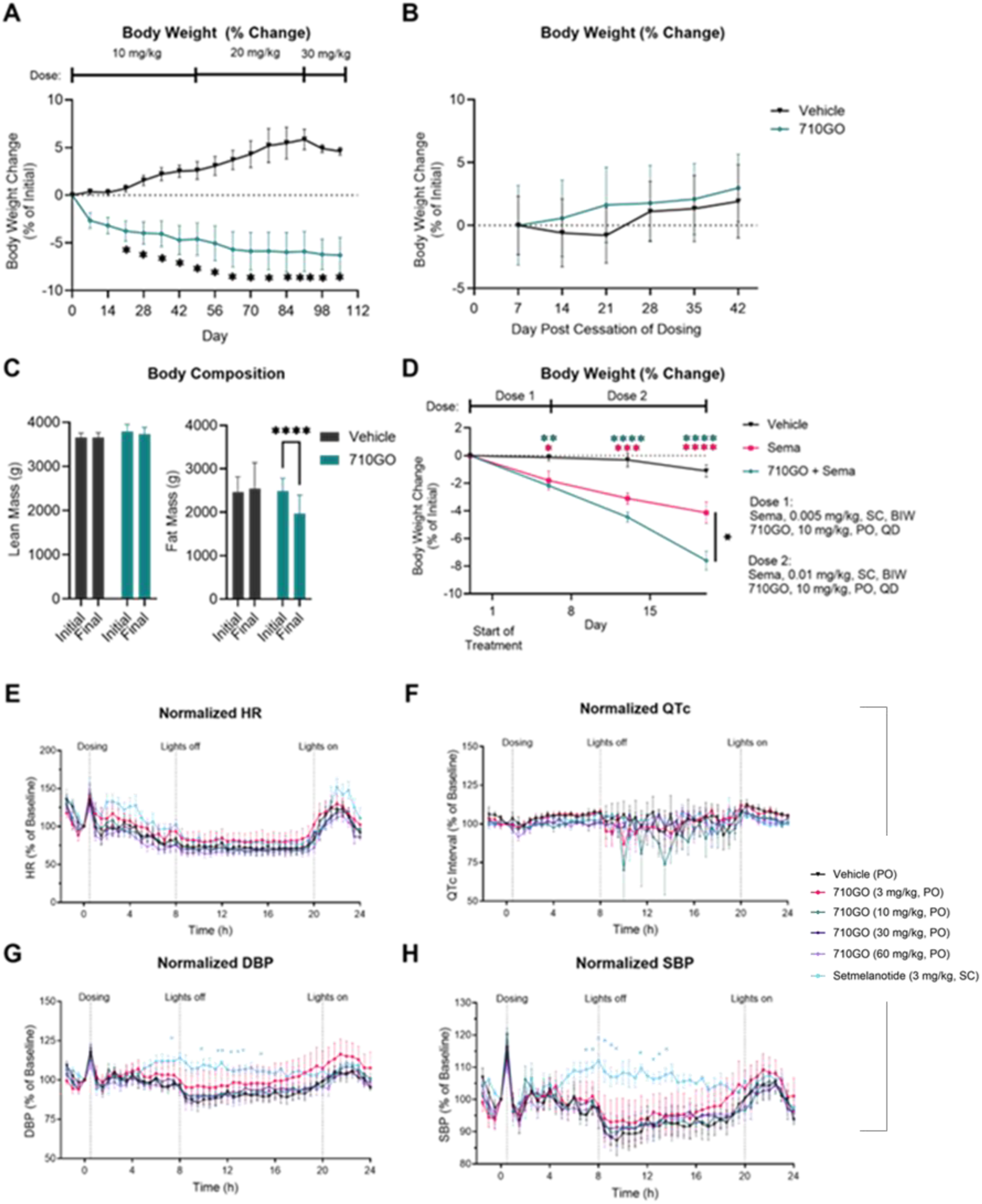
Effects of 710GO on body weight, mass composition, weight regain, GLP-1 administration, and cardiac parameters in DIO NHPs. **A)** Body weight change in response to oral administration of 710GO (green diamond, n=10) compared to saline vehicle (inverted black triangle, n=4). **B)** Change in body weight from initial baseline after dose cessation of 710GO (n=10) and vehicle (n=4) over six weeks. **C)** Changes in fat mass and lean mass in response to oral administration of 710GO (n=10) and saline vehicle (n=4). **D)** Change in body weight in response to saline vehicle (inverted black triangle, n=8) and semaglutide (Sema, pink square, n=3), and combination of oral 710GO and subcutaneous Sema (green diamond, n=4). The Sema dose was escalated from 0.005 mg/kg SC BIW to 0.01 mg/kg SC BIW on Day 8 in both the individual treatment group and in combination with 710GO (10 mg/kg, PO, QD). **E-H)** Effects of 710GO (n=5 per group) at 3 mg/kg (pink circle), 10 mg/kg (green diamond), 30 mg/kg (dark purple square), and 60 mg/kg (light purple diamond), PO, QD, and setmelanotide (n=3, 3 mg/kg, SC, QD, light blue circle) on normalized heart rate (E), QT interval (F), diastolic blood pressure (G), and systolic blood pressure (H). Data are presented as mean ± SEM. Statistical significance in (E,F) using mixed-effects analysis, followed by Dunnett’s multiple comparison test (*p < 0.05).

Given the reported concerns of GLP-1 therapy, we specifically evaluated 710GO for weight rebound, loss of lean mass, and GI side effects. ^2,30,31^ After dosing concluded in our 15-week study, DIO NHPs were monitored for an additional six weeks. Two weeks after cessation of treatment, NHPs treated with 710GO lost an additional 0.39% of their initial body weight. By six weeks after treatment cessation, they had only regained 34.2% of the total weight lost during treatment. This slope of weight regain was not significantly different when compared to animals receiving placebo (**Fig. 4B**). In a separate 35-day study, 710GO-treated DIO NHPs lost an average of 518.1 ± 132.7g of fat mass, corresponding to 90.1% of the total mass lost (**Fig. 4C**). Notably, 710GO did not significantly decrease lean mass in DIO NHPs, with lean mass only constituting 9.93% of the total mass lost (**Fig. 4C**). 710GO-treated animals appeared healthy, with no observed vomiting or diarrhea. We then evaluated 710GO as an adjuvant to semaglutide for weight loss in DIO NHPs over 19 days. Semaglutide alone decreased body weight by 4.1 ± 0.8% compared to 7.6 ± 0.7% when co-administered with 710GO (**Fig. 4D, Supplementary Fig. 6D**). Not only was the combination treatment significantly more effective than semaglutide alone, it also did not induce vomiting or GI distress (**Fig. 4D**).

Melanocortin agonists have previously been limited by cardiac side effects.^32^ To investigate the effect of 710GO on cardiac parameters, increasing doses were orally administered to NHPs implanted with telemetry devices. Over a 24-hour monitoring period, 710GO did not induce changes in cardiac parameters, including systolic blood pressure (SBP), diastolic blood pressure (DBP), heart rate (HR), and corrected QT interval (QTc) at doses up to 60 mg/kg (**Fig. 4E-H**). In comparison, setmelanotide (an FDA-approved melanocortin agonist) significantly increased nighttime DBP and SBP and decreased body temperature several hours after a 3 mg/kg subcutaneous dose (**Supplementary Fig. 11**). Both 710GO (60 mg/kg, PO) and setmelanotide (3 mg/kg, SC) were dosed six times higher than their previously established efficacious dose in NHPs.^33^ Additionally, 710GO did not result in hERG inhibition (**Supplementary Fig. 12**). These results demonstrate that 710GO safely modulates the melanocortin system without increasing sympathetic tone and may also have a greater therapeutic window than setmelanotide.

The respective contributions of MC3R and MC4R to energy homeostasis have long remained elusive. Historically, MC4R was viewed as the primary mediator of food intake and weight regulation, while MC3R was considered ancillary.^16–20^ While humans with MC4R LoF mutations display frank hyperphagia and obesity, MC3R LoF variants typically present with a mild overweight phenotype.^22,34^ Yet, recent work suggests that MC3R functions as a rheostat, which modulates downstream melanocortin tone based on peripheral cues of energetic state.^14^ To dissect this interplay, we evaluated selective MC3R, MC4R, and dual MC3R/MC4R agonists in NHPs under both acute and chronic metabolic perturbations.

In metabolically normal, lean animals under homeostatic conditions, selective activation of either receptor had little effect on suppressing feeding, while dual agonism produced a modest reduction of food intake. However, during acute fasting, MC3R agonism alone markedly suppressed refeeding behavior in lean animals, whereas MC4R agonism alone did not. Combined MC3R/MC4R activation further enhanced this anorectic effect, indicating that MC3R plays a dominant role in responding to a transient energy deficit and that dual activation amplifies this response.

In obese NHPs undergoing subchronic dosing, receptor contributions diverged: selective MC4R agonism reduced food intake and body weight, consistent with its established role in long-term energy regulation, while dual MC3R/MC4R agonism produced greater and more durable reductions across both metrics. In contrast, selective MC3R agonism did not decrease food intake and was paradoxically associated with modest weight gain. This finding aligns with previous studies, where *Mc3rKO* animals exposed to obesogenic challenges vastly overshoot an appropriate set point compared to wildtype controls, indicating that MC3R is necessary for setting an upper limit on metabolic regulation.^14^

These findings delineate a potential functional division of labor within the melanocortin system. MC3R acts as a contextual modulator, tuning the magnitude of the response in line with metabolic state, while MC4R drives sustained changes in feeding and body weight. Together, they operate as an integrated network, coordinating adaptive and chronic control of body weight. Thus, modulating MC3R is necessary to adjust the boundary and allow for more profound weight gain or loss through MC4R activation above the previously established metabolic set point.

One additional hypothesis emerging from these data is that obesity may blunt responsiveness to tonic α-MSH–mediated food inhibition, rendering melanocortin signaling partially desensitized.^35,36^ MC4R is known to have high basal activity, which may contribute to the observed reduction in food intake when selectively agonizing MC3R in lean fasted animals.^37^ However, in obese NHPs, a loss of high constituent MC4R tone renders selective MC3R agonism insufficient to lower food intake. Under such conditions, supraphysiological MC3R and MC4R agonism may help restore this lost sensitivity by amplifying melanocortin tone or re-establishing inhibitory thresholds, thereby enabling more effective MC4R signaling.

Our findings also contextualize prior clinical programs. Early MC4R-selective agonists achieved only limited efficacy, even after prolonged treatment.^38,39^ More recent compounds, such as setmelanotide and bivamelagon, retain some potency on MC3R and produced greater weight loss, supporting the hypothesis that MC3R engagement enhances MC4R-driven outcomes.^40,41^ However, limitations in dosing due to potential off-target adverse effects, including increased sympathetic tone, likely prevented full receptor activation and clinically-relevant weight loss.^42^ To overcome these limitations, we developed 710GO, an equipotent dual MC3R/MC4R agonist. High potency at both receptors allows for efficacy at lower exposures, a C-terminal extension minimizes off-target interactions, and non-canonical amino acids enhance physiological stability.^43,44^ These design features collectively enable precise central melanocortin engagement with improved efficacy and tolerability.

While promising, these studies were conducted with modest sample sizes and showed large interindividual variability. The compounds used in this study are selective relative to MC3R and MC4R; however, they also have varying activity on MC1R and MC5R and their contributions cannot be ruled out. Across studies, rodent body weight was always tightly correlated to changes in food intake, whereas in NHPs, the two metrics diverged. These differences may reflect important evolutionary and ecological pressures that led to more nuanced mechanisms for metabolic control, such as alterations in energy partitioning. These observations warrant further investigation. Regardless, dual agonism consistently resulted in the greatest reduction in both food intake and body weight, irrespective of species. Future work, including a first-in-human phase I trial, will evaluate the translational potential of MC3R/MC4R dual activation.

Collectively, these findings position MC3R/MC4R dual activation as a novel mechanism that re-establishes the melanocortin system as a promising target for obesity treatment. Dual MC3R/MC4R agonism recapitulates the normal physiological integration required for effective and sustainable weight regulation. By engaging both receptors, 710GO achieves efficacy at lower exposures and thus a broader safety margin than prior MC4R-selective compounds. Moreover, the innate cooperation between melanocortin and incretin systems lends itself to combination therapy with lower doses, greater efficacy, and reduced side effects. Additionally, we observe minimal weight rebound and preferential loss of fat mass relative to lean mass with orally-administered 710GO, which could provide a mechanism of durable, healthier weight loss. After decades of limited success in melanocortin-based obesity pharmacotherapy, these findings offer a renewed path forward.

## Methods

### Compound procurement

Peptides and small molecules were either synthesized at WuXi AppTec or purchased from commercial vendors.

### Cyclic AMP assays

CHO-K1 cells expressing recombinant human receptor grown prior to the test in media without antibiotic were detached by gentle flushing with PBS-EDTA (5 mM EDTA), recovered by centrifugation and resuspended in assay buffer (KRH: 5 mM KCl, 1.25 mM MgSO_4_, 124 mM NaCl, 25 mM HEPES, 13.3 mM Glucose, 1.25 mM KH_2_PO_4_, 1.45 mM CaCl_2_, 0.5 g/l BSA, supplemented with 1 mM IBMX or 25 mM Rolipram). Dose response curves were performed in parallel with the reference compounds. For agonists test (384 wells): 5 mL of cells were mixed with 5 mL of test compound at concentrations from 0.0001 nM to 1000 nM and then incubated 30 minutes at room temperature. After addition of the lysis buffer containing cAMP-d2 and anti-cAMP cryptate detection reagents, plates were incubated 1 hour at room temperature, and fluorescence ratios were measured according to HTRF kit specifications. For antagonists test (384 wells): 5 mL of cells were mixed with 5 mL of test compound at concentrations from 0.0001 nM to 1000 nM and reference agonist NDP-α-MSH for a final concentration corresponding to the historical EC_80_. Plates were then incubated for 30 minutes at room temperature. After addition of the lysis buffer containing cAMP-d2 and anti-cAMP cryptate detection reagents, plates were incubated 1 hour at room temperature, and fluorescence ratios were measured according to HTRF kit specifications.

### Cynomolgus macaque husbandry

All protocols and any amendments or procedures involving the care or use of animals on this study were reviewed and approved by WuXi AppTec Institutional Animal Care and Use Committee (IACUC) prior to the initiation of such procedures (IACUC No. 24043001). Animals were individually housed in the stainless-steel mesh cages in accordance with local standard and Testing Facility SOPs. Lean animals were supplied with a certified Monkey Diet twice each day, except for overnight fasting for acute refeeding experiments where specified. DIO-NHPs were fed a high fat diet for the entirety of the study, including standard chow (3.5 kcal/g) as the morning meal, high-fat chow (4.31 kcal/g) as the afternoon meal. In addition, animals received fruit (0.52 kcal/g) daily as nutritional enrichment. Facilities were kept at a humidity between 40% to 70% at temperatures ranging from 16 to 26°C with at least 8 air changes/hour. An electronic time-controlled lighting system was used to provide a 12-hour light/12-hour dark cycle. The animals were supplied with fresh reverse-osmosis (RO) water using an automated watering system. Veterinary care was available and provided to the animals throughout the course of the study. Any decisions about removing animals from studies for health reasons were made at the judgement of veterinary staff.

### Selective and combination agonism acute feeding studies in cynomolgus macaques

Fifteen non-naïve adult, male cynomolgus macaques between the ages of four and six years old were weighed (starting body weight 3-6 kg) and baseline food intake was measured for 6 days prior to the start of dosing. Animals were then grouped such that the average food intake, age and body weight in each group was roughly equivalent. The study was performed in a within-subjects design where each animal acts as its own control by receiving a single administration of each dose level, including vehicle. Macaques were given SC, QD, either vehicle (300 mM mannitol in sWFI), EBMC-40-187 (an MC3R-selective agonist), EBMC-03-47 (an MC4R-selective agonist), or combination EBMC-40-187 and EBMC-03-47 (n=3, per dose group, except where one animal that was removed from the *ad libitum* combination dose group based on a ROUT outlier analysis, Q=1%). In both the EBMC-40-187 and the combination dosing group, EBMC-40-187 was administered at 8 and 15 mg/kg. In both the EBMC-03-47 and the combination dosing group, EBMC-03-47 was administered at 3 and 6 mg/kg. Food intake was monitored for 24 hours post-dose. Macaques were regularly observed via cage-side observation for the duration of the study.

### MC3R selective agonism, MC4R selective agonism, and MC3R + MC4R co-agonism sub-chronic feeding study in cynomolgus macaques

Twenty-five naïve adult (ages 9-20 years old), male diet-induced obese cynomolgus macaques with starting body weights >8.5 kg were weighed and baseline food intake was measured for 6 days prior to the start of dosing. Animals were then grouped such that the average food intake, age and body weight in each group was roughly equivalent, with extra animals being released back into the colony. Macaques were given SC, QD, either vehicle (300 mM mannitol in sWFI), EBMC-40-187 (an MC3R-selective agonist), EBMC-03-47 (an MC4R-selective agonist), or combination EBMC-40-187 and EBMC-03-47 for 24 days (n=5). In both the EBMC-40-187 and the combination dosing group, EBMC-40-187 was administered at 4 mg/kg for days 1-10 and 8 mg/kg for days 11-24. In both the EBMC-03-47 and the combination dosing group, EBMC-03-47 was administered at 1.5 mg/kg for days 1-10 and 3 mg/kg for days 11-24. Food intake was monitored by meal. Body weight was measured weekly and on day 24 (the final day of dosing). Animals were fasted overnight for the collection of blood for clinical chemistry before the start of the study and every two weeks until the end of dosing. Macaques were monitored by regular cage-side observation for the duration of the study.

### MC3/4R Co-Agonism (710GO) and MC4R agonism/MC3R partial antagonism (707GO) acute feeding studies in cynomolgus macaques

Six non-naïve adult, male cynomolgus macaques between the ages of 10-25 years old were weighed (starting body weight >8 kg) and baseline food intake was measured for 6 days prior to the start of dosing. Animals were then grouped such that the average food intake, age and body weight in each group was roughly equivalent. The study was performed in a within-subjects design where each animal acts as its own control by receiving a single administration of each dose level, including vehicle. Macaques were given SC, QD, either vehicle (saline), 710GO (3 mg/kg, SC, or 707GO (n=2, per dose group). Food intake was monitored for 72 hours post-dose. Macaques were regularly observed via cage-side observation for the duration of the study.

### MC3/4R Co-Agonism (710GO) and MC4R agonism/MC3R partial antagonism (707GO) sub-chronic feeding study in cynomolgus macaques

Forty non-naïve adult (ages 6-22 years old) male, diet-induced-obese cynomolgus macaques with starting body weight >8.5 kg were weighed and food intake was measured for 6 days prior to the start of the study. Animals were then grouped such that the average food intake, age and body weight in each group was roughly equivalent, with extra animals being released back into the colony. They were then administered either vehicle (saline, n=16), 710GO (n=15), or 707GO (n=8). For the 710GO and 707GO groups, macaques were administered 10 mg/kg PO, QD for days 1-8, 20 mg/kg PO, QD for days 9-14, and 15 mg/kg PO, BID for days 15-35. Food intake was monitored by meal. Body weight was measured weekly. Animals were fasted overnight for the collection of blood for clinical chemistry before the start of the study and every two weeks until the end of dosing. Macaques were monitored by regular cage-side observation for the duration of the study.

### Chronic dual-agonist (710GO) feeding study

Fourteen adult (ages 10-22) male, diet-induced-obese cynomolgus macaques with starting body weights >8 kg, were administered either vehicle (n=4) or 710GO by oral gavage. The 710GO treatment group was split into two cohorts. The first 710GO cohort (n=4) received 10 mg/kg for days 1-91. The second 710GO cohort (n=6) received 10 mg/kg for days 1-51, 20 mg/kg for days 52-91, and 30 mg/kg for days 92-105. The 710GO results from the study are described as the combined data from both 710GO cohorts. Food intake was monitored by meal. Body weight was measured weekly, and weight rebound was assessed for 6 weeks after cessation of the treatment phase. Blood labs were taken at the start of the study and every 2 weeks until the end of dosing. One animal was dropped from the saline group due to nonspecific illness. Body composition was measured by MRI at the beginning and end of the treatment phase of the second 710GO cohort.

### Subchronic combination 710GO + GLP-1 feeding study in cynomolgus macaques

Sixteen adult non-naïve male, DIO (>8 kg) NHPs, ages 6-22, were treated with vehicle (saline, n=8), semaglutide (n=3, one animal was removed from the study for nonspecific illness) or combination 710GO and semaglutide (n=4). Semaglutide was administered at a dose of 0.005 mg/kg SC, BIW for week 1, and 0.01 mg/kg SC, BIW for weeks 2-3. 710GO was administered 10 mg/kg PO, QD for days 1-19. The combination group was treated with semaglutide (0.005 mg/kg SC, BIW) and 710GO (10 mg/kg, PO, QD) for week 1 and semaglutide (0.01 mg/kg SC, BIW) and 710GO (10 mg/kg, PO, QD) for weeks 2-3. Food intake was monitored by meal. Body weight was measured weekly. Blood labs were taken at the start of the study and every 2 weeks until the end of dosing. Macaques were monitored by regular cage-side observation for the duration of the study.

### Cardiac safety studies of 710GO in cynomolgus macaques

Five non-naïve adult male cynomolgus macaques between 6-9 years old, with starting body weights between 7-9 kg, were surgically implanted with telemetry devices to monitor body temperature and cardiac parameters including SBP, DBP, HR, and QTc. Each animal (n=5) received vehicle (sWFI) and 710GO at doses of 3, 10, 30, and 60 mg/kg, PO, in a Latin square design. Setmelanotide was dosed 3 mg/kg, SC (n=3). Animals were dosed in the morning; body temperature and cardiac parameters were measured every 30 min over a 24-hour monitoring period.

### Statistical Methods

Statistical analyses were performed using GraphPad Prism (v10.6.1). Sample sizes (n) refer to the number of animals in each group. Data are shown as mean ± SEM. For comparisons among more than two groups, one- or two-way ANOVA or mixed-effects analyses followed by either Dunnett’s (when comparing only to vehicle group), uncorrected Fisher’s LSD, or Tukey’s multiple-comparison test were applied. A p-value < 0.05 was considered statistically significant. Specific n, test used and p-value are reported in all figure legends.

## Data Availability

All data are available upon reasonable request made to DLM.

## Acknowledgments

We thank Andres Arango and Sanjay L Kumar for reviewing and providing feedback on the manuscript. We also thank the staff members of WuXi AppTec and EuroScreenFast who contributed to animal care and research activities.

