## Supplemental for "Dual Activation of MC3R and MC4R Drives Weight Loss and Reduces Food Intake in Obese Primates"

### Supplementary Figures:

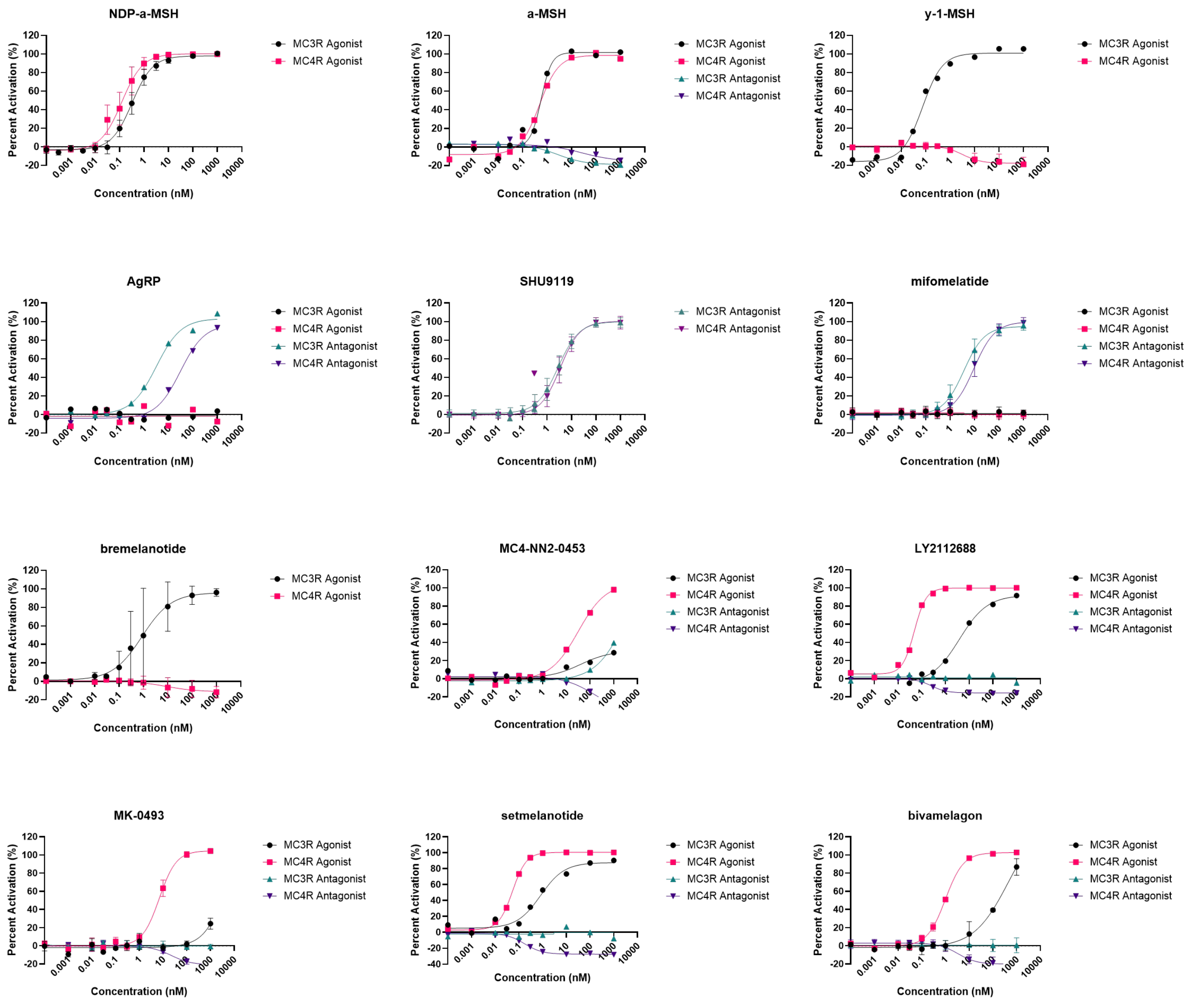


**Supplementary Fig. 1 |** Pharmacological characterization of melanocortin ligands against human MC3R and human MC4R expressed in CHO-K1 cells by cAMP signaling. Percent activation is relative to NDP-a-MSH for agonist mode and SHU9119 for antagonist mode.

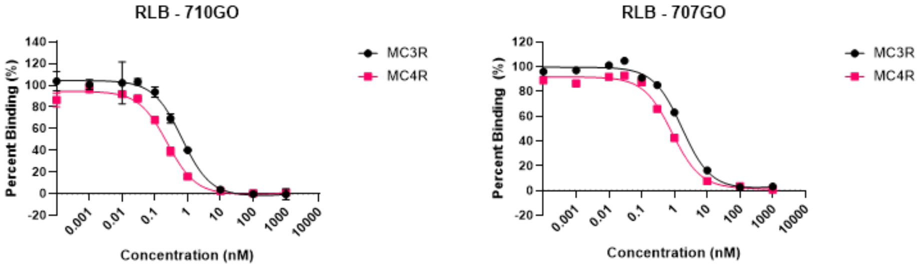


| Compound ID | MC3R $IC_{50}$ (nM) | MC4R $IC_{50}$ (nM) |
|---|---|---|
| 710GO | 0.64 | 0.81 |
| 707GO | 1.66 | 0.23 |

**Supplementary Fig. 2 |** Radiolabeled binding (RLB) competition activity on recombinant human MC3R and human MC4R expressed in CHO-K1 cells. Percent of [125I]-NDP-a-MSH bound after addition of 710GO or 707GO.

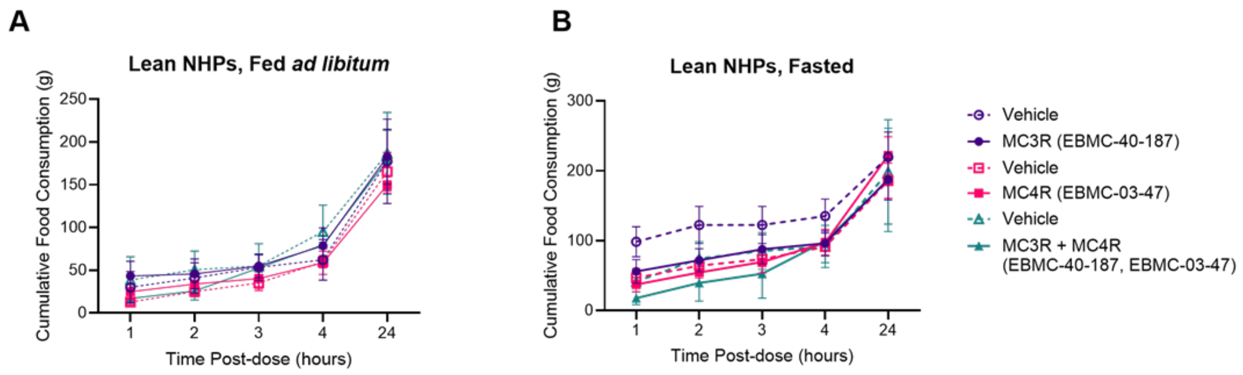


**Supplementary Fig. 3 |** Individual and combined effects of MC3R and MC4R agonism on cumulative food intake in lean NHPs (n=3). A,B) Cumulative food intake (g) in lean, NHPs fed *ad libitum* (A) and lean, fasted NHPs (B) after subcutaneous injection of vehicle (open circle, square, and triangle, 300 mM mannitol in sWFI), selective MC4R agonist EBMC-03-47 (pink filled square, 3 mg/kg), selective MC3R agonist EBMC-40-187 (purple filled circle, 8 mg/kg), and co-administration of the two agonist compounds (green filled triangle, 3 mg/kg MC4R agonist, 8 mg/kg MC3R agonist) over a 24-hour monitoring period.

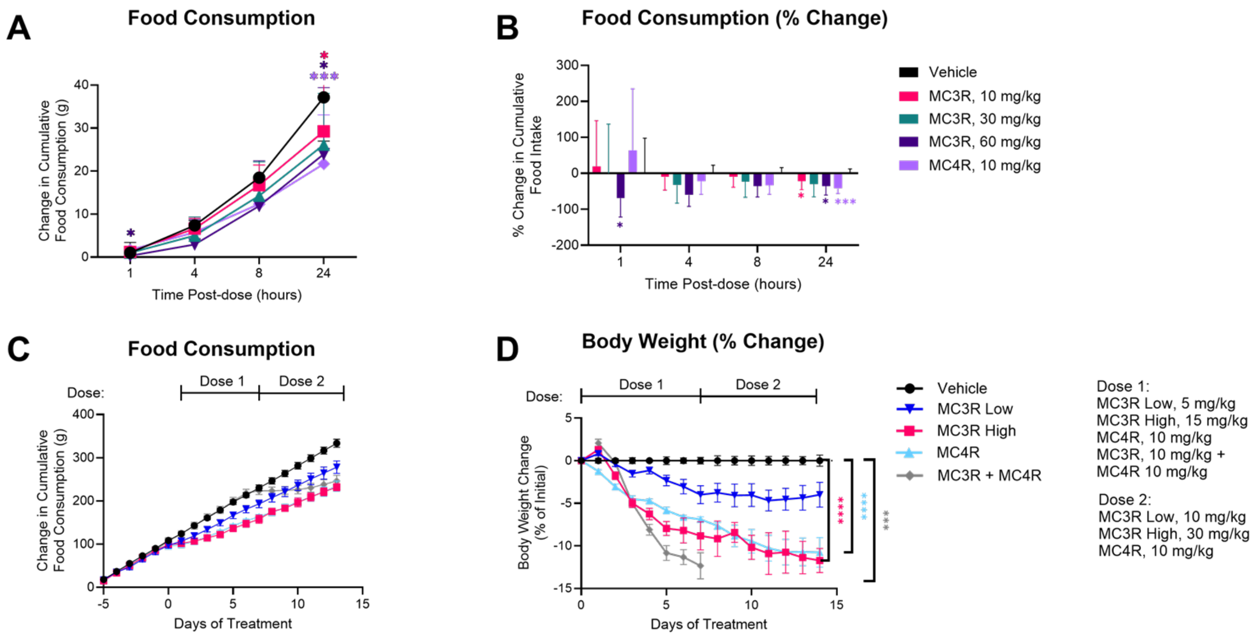


**Supplementary Fig. 4 |** Agonism of MC3R or MC4R is sufficient to reduce food intake and body weight in DIO rats (n=6, per test article dose group). **A)** Change in cumulative food intake (g) after subcutaneous injection of vehicle (black circle, 300 mM mannitol in sWFI, n=12), selective MC4R agonist EBMC-03-47 (10 mg/kg, light purple diamond) and selective MC3R agonist EBMC-40-187 (10 mg/kg pink square, 30 mg/kg green triangle, 60 mg/kg dark purple inverted triangle) over a 24-hour monitoring period. **B)** Percent change in cumulative food between test compounds and vehicle. **C,D)** Individual and combined effects of MC3R and MC4R agonism on cumulative food intake (C) and body weight (D) in DIO rats over 14 days of treatment. Test articles including vehicle (black square, 300 mM mannitol), selective MC4R agonist EBMC-03-47 (light blue triangle), selective MC3R agonist EBMC-40-187 (blue inverted triangle/pink square), and co-administration of MC3R/MC4R agonists (grey diamond) were administered once daily by subcutaneous injection. EBMC-40-187 doses were increased from 5 mg/kg to 10 mg/kg (low) and 15 mg/kg to 30 mg/kg (high) on Day 7. Combination treatment began on Day 7 and was aligned with the Day 0 start of other groups. The combination group was administered EBMC-40-187 (30 mg/kg, Day 0-1; 10 mg/kg Day 2-7) and EBMC-03-47 (10 mg/kg, Day 0-7). Cumulative food intake was monitored for five days prior to administration of first dose. Statistical significance in (A,B) vs vehicle using Two-Way ANOVA and (C,D) using One-Way ANOVA, both followed by Dunnett's multiple comparison test (*$p < 0.05$, ***$p<0.001$, ****$p<0.0001$).

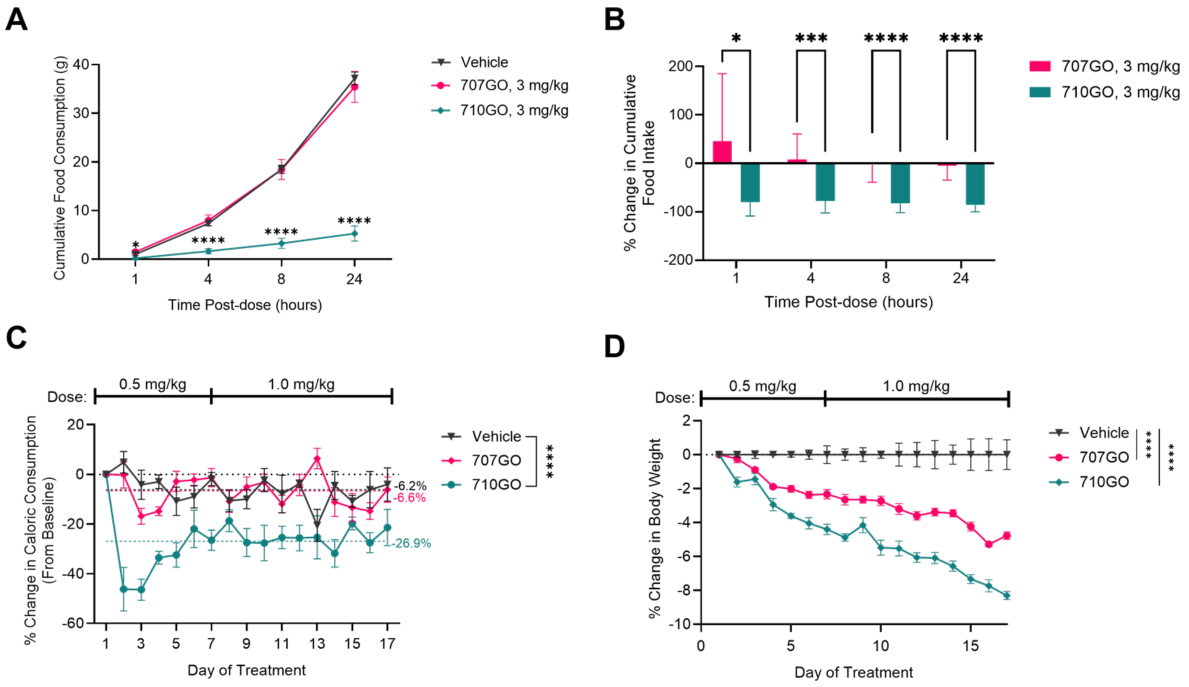


**Supplementary Fig. 5 |** Dual agonism of MC3R and MC4R by 710GO potently reduces food intake and body weight in DIO rats. **A-B)** Cumulative (A) and percent change in cumulative (B) food intake in DIO rats during a 24-hour acute feeding study induced by 710GO (MC3R/MC4R dual agonist, 3 mg/kg, green diamond, n=12) and 707GO (partial MC3R antagonist/MC4R agonist, 3 mg/kg, pink circle, n=12) compared to vehicle (300 mM mannitol in sWFI, black inverted triangle, n=12). **C-D)** Percent change in cumulative caloric consumption (C) and in body weight (D) in DIO rats during a sub-chronic 17-day study induced by daily administration of 710GO (green diamond, n=5) and 707GO (pink circle, n=5) compared to vehicle (saline, black inverted triangle, n=4). In sub-chronic studies, 710GO and 707GO were administered subcutaneously at doses of 0.5 mg/kg for days 1-7 and 1.0 mg/kg for days 8-17. Statistical significance in (A,B) vs vehicle using One-Way ANOVA and (C,D) using Two-Way ANOVA, both followed by Dunnett's multiple comparison test (*p < 0.05, ***p<0.001, ****p<0.0001).

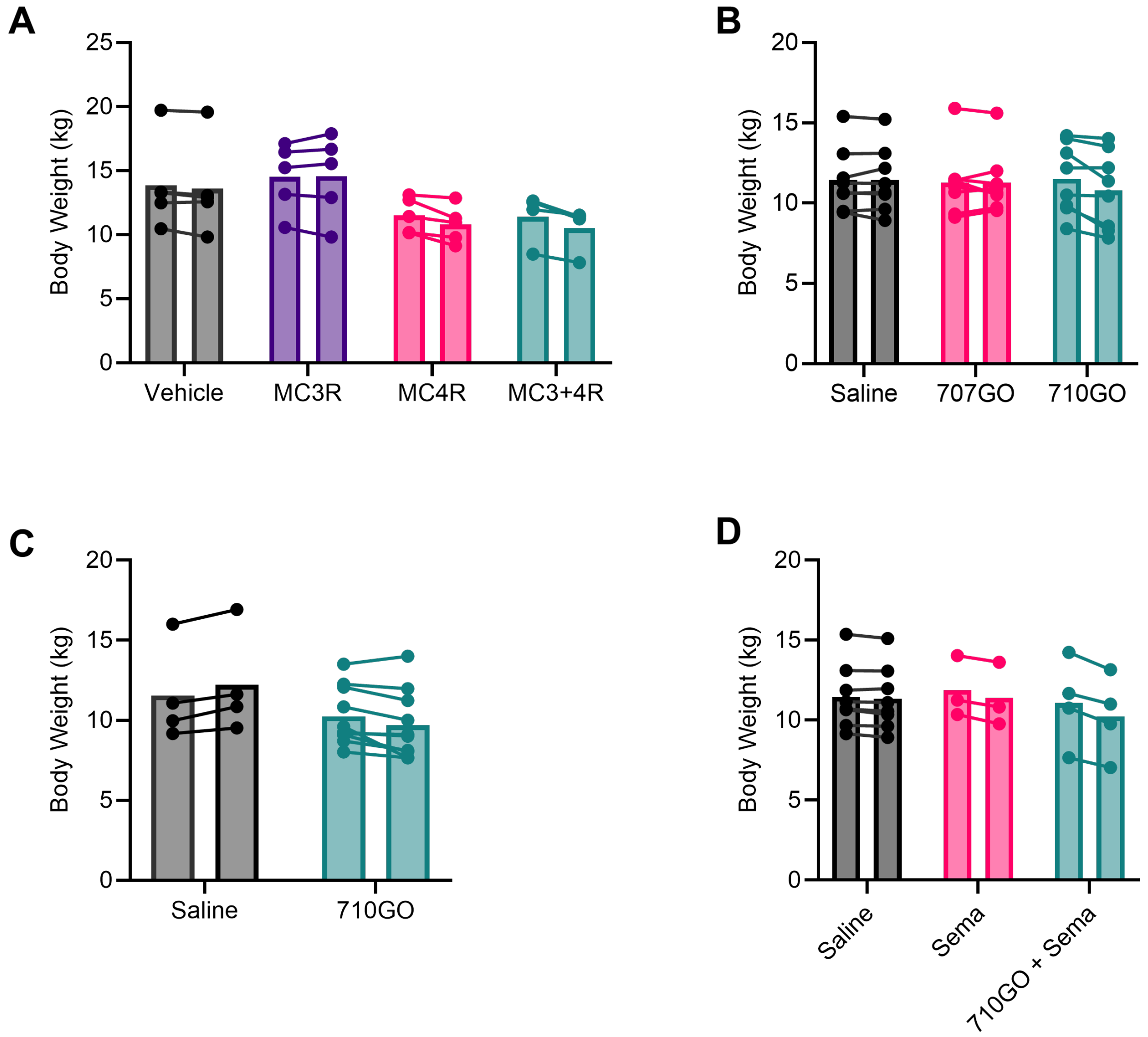


**Supplementary Fig. 6 |** Body weight measurements for NHPs receiving peptide therapeutics. Dosing information for each can be found in main text figures referenced below. **A)** Raw body weight in DIO male NHPs on days 1 and 25, as described in Figure 2D. Animals were administered vehicle (300 mM mannitol in sWFI, black) or test compound by subcutaneous injection (n=5, per dose group; n=4 for co-agonism group). EBMC-40-187 (purple) is a full MC3R agonist, EBMC-03-47 (pink) is a full MC4R agonist, and the two peptides were co-administered for MC3R + MC4R co-agonism (green). **B)** Raw body weight in NHPs during repeated dosing of either vehicle (saline, black), 707GO (pink), or 710GO (green) on days 1 and 36 (n=8 per group), as described in Figure 3D. **C)** Raw body weight on day 1 and 91 in response to oral administration of 710GO (green, n=10) compared to vehicle (saline, black, n=4), as described in Figure 4A. **D)** Raw body weight on day 1 and 17 in response to vehicle (saline, black, n=8), semaglutide (Sema, pink, n=3), and combination of oral 710GO and subcutaneous Sema (green, n=4), as described in Figure 4D.

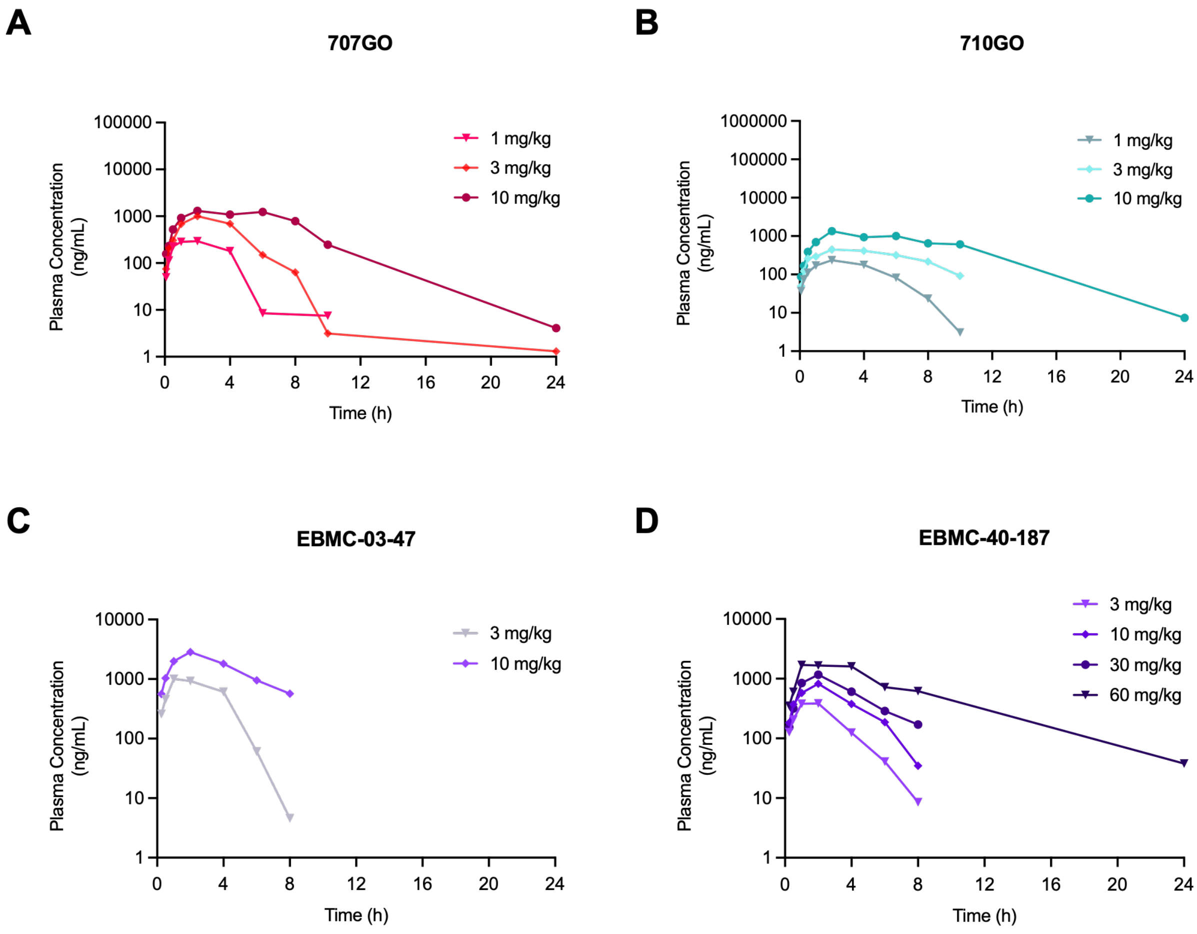


**Supplementary Fig. 7 |** Plasma concentration of melanocortin agonist peptides after subcutaneous injection in rats (n=3, per dose group) over a 24-hour monitoring period. Test articles 707GO (**A**), 710GO (**B**), EBMC-03-47 (**C**), and EBMC-40-187 (**D**) were administered at various concentrations including 1 mg/kg (circles), 3 mg/kg (triangles), 10 mg/kg (squares), 30 mg/kg (inverted triangles), and 60 mg/kg (diamonds). Not all doses were investigated for each test article.

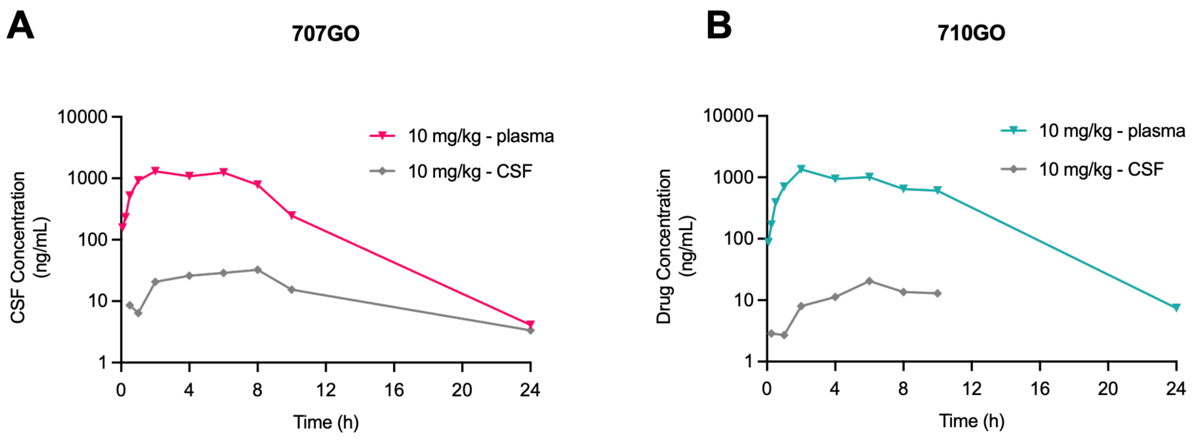


**Supplementary Fig. 8 |** Concentration of 707GO (**A**) and 710GO (**B**) in plasma (inverted triangles) and cerebrospinal fluid (diamonds) after subcutaneous injection at a dose of 10 mg/kg in rats (n=3, per dose group) over a 24-hour monitoring period.

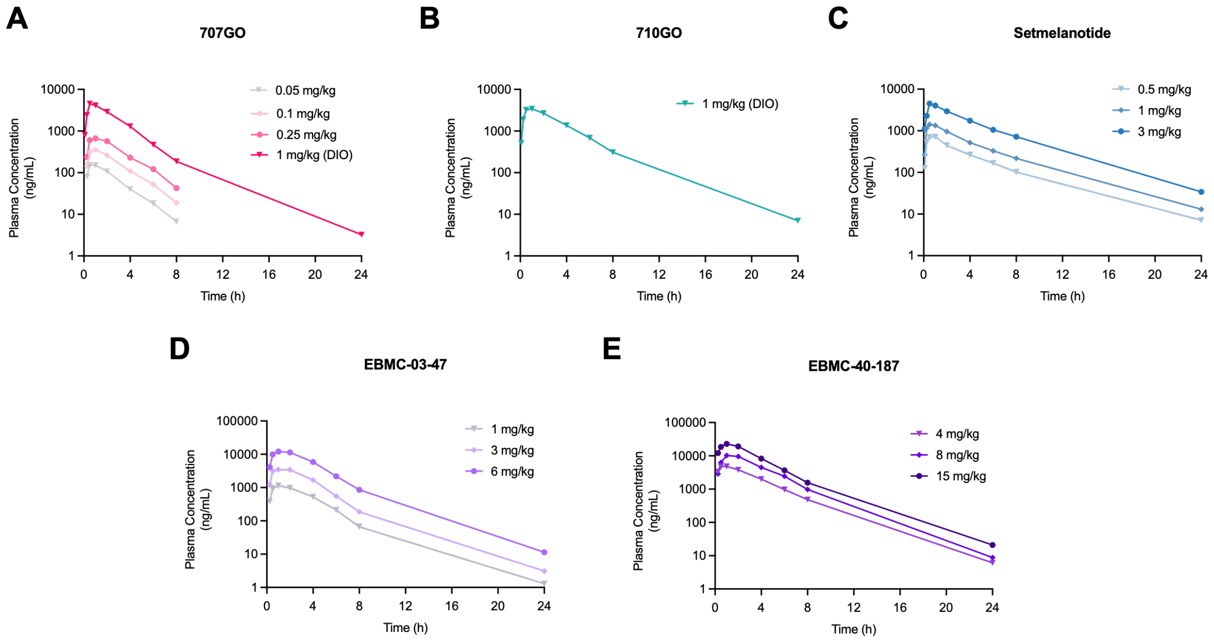


**Supplementary Fig. 9 |** Plasma concentration of melanocortin agonist peptides after subcutaneous injection in lean NHPs (n=3, per dose group) and/or DIO NHPs (n=2, per dose group) over a 24-hour monitoring period. **A**) Plasma concentration of 707GO after administration at doses of 0.05 mg/kg (grey inverted triangle), 0.1 mg/kg (light pink diamond), and 0.25 mg/kg (medium pink circle) in lean NHPs and 1 mg/kg (hot pink inverted triangle) in DIO NHPs. **B**) Plasma concentration of 710GO after administration at a dose of 1 mg/kg (green inverted triangle) in DIO NHPs. **C**) Plasma concentration of setmelanotide after administration at doses of 0.5 mg/kg (light blue inverted triangle), 1 mg/kg (medium blue diamond), and 3 mg/kg (dark blue circle) in lean NHPs. **D**) Plasma concentration of EBMC-03-47 after administration at doses of 1 mg/kg (grey inverted triangle), 3 mg/kg (light purple diamond), and 6 mg/kg (dark purple circle) in lean NHPs. **E**) Plasma concentration of 707GO after administration at doses of 4 mg/kg (light purple inverted triangle), 8 mg/kg (medium purple diamond), and 15 mg/kg (dark purple circle) in lean NHPs.

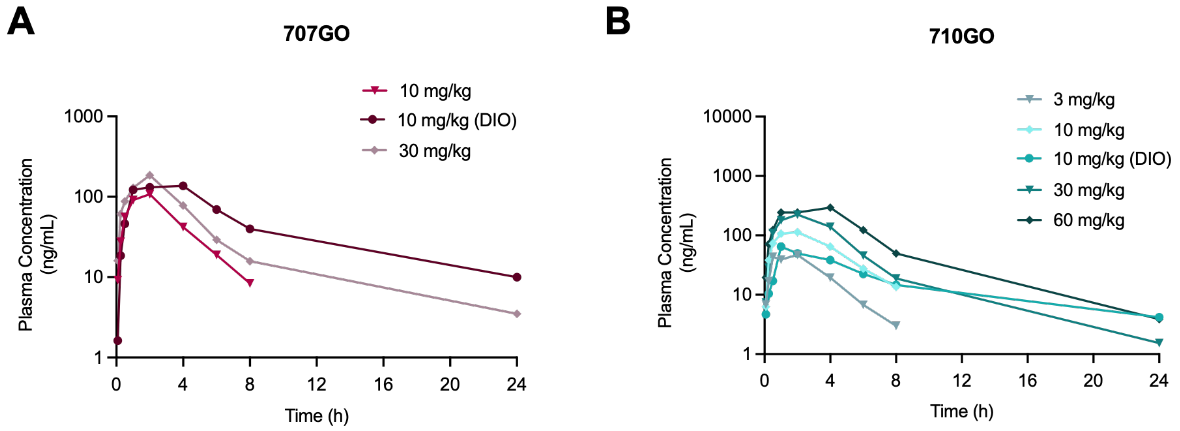


**Supplementary Fig. 10 |** Plasma concentrations of 707GO and 710GO after administration by oral gavage in NHPs over a 24-hour monitoring period. A) Plasma concentration of 707GO after oral administration at doses of 10 mg/kg (red inverted triangle) and 30 mg/kg (maroon circle) in lean NHPs (n=3, per dose group) and 10 mg/kg (grey diamond) in DIO NHPs (n=4, per dose group). B) Plasma concentration of 710GO after oral administration at doses of 3 mg/kg (light blue inverted triangle), 10 mg/kg (cyan diamond), 30 mg/kg (dark teal inverted triangle), and 60 mg/kg (dark green diamond) in lean NHPs (n=3, per dose group) and 10 mg/kg (light teal circle) in DIO NHPs (n=4, per dose group).

**Supplementary Table 1 |** Pharmacokinetic parameters in plasma of melanocortin agonist peptides after subcutaneous injection in rats at various dose concentrations. All data are presented as mean ± SD.

| | | PK Parameters in Sprague-Dawley Rats | | | | |
|---|---|---|---|---|---|---|
| **Compound** | **Subcutaneous Injection Dose** | **$C_{max}$ ± SD (ng/mL)** | **$T_{max}$ ± SD (h)** | **$T_{1/2}$ ± SD (h)** | **$AUC_{0\text{-}last}$ ± SD (ng*h/mL)** | **$AUC_{0\text{-}inf}$ ± SD (ng*h/mL)** |
| **707GO** | **1 mg/kg (n=3)** | 297 ± 19 | 1.67 ± 0.58 | 1.03 ± 0.48 | 1066 ± 59 | 1077 ± 58 |
| | **3 mg/kg (n=3)** | 1011 ± 146 | 2.00 ± 0.00 | 1.93 ± 1.06 | 3877 ± 225 | 3881 ± 227 |
| | **10 mg/kg (n=3)** | 1384 | 2.00 | 2.17 | 10475 | 10488 |
| **710GO** | **1 mg/kg (n=3)** | 238 | 1.00 | 2.59 | 1090 | 1124 |
| | **3 mg/kg (n=3)** | 448 | 0.75 | 2.84 | 3002 | 3450 |
| | **10 mg/kg (n=3)** | 1366 | 2.00 | 2.78 | 10731 | 10781 |
| **EBMC-03-47** | **3 mg/kg (n=3)** | 1005 ± 33 | 1.00 ± 0.00 | 0.58 ± 0.10 | 3552 ± 350 | 3558 ± 356 |
| | **10 mg/kg (n=3)** | 2863 | 2.00 | 2.94 | 12560 | 15643 |
| **EBMC-40-187** | **3 mg/kg (n=3)** | 426 ± 150 | 1.33 ± 0.58 | 1.11 ± 0.31 | 1292 ± 356 | 1308 ± 344 |
| | **10 mg/kg (n=3)** | 830 | 2.00 | 1.29 | 3007 | 3091 |
| | **30 mg/kg (n=3)** | 1178 ± 212 | 2.00 ± 0.00 | 2.33 ± 0.70 | 4481 ± 979 | 5124 ± 1143 |
| | **60 mg/kg (n=3)** | 1722 | 2.00 | 4.16 | 13028 | 13272 |

**Supplementary Table 2 |** Pharmacokinetic parameters in plasma of melanocortin agonist peptides after subcutaneous injection in NHPs at various dose concentrations. All data are presented as mean ± SD.

| | | PK Parameters in Cynomolgus Macaques | | | | |
|---|---|---|---|---|---|---|
| **Compound** | **Subcutaneous Injection Dose** | **$C_{max}$ ± SD (ng/mL)** | **$T_{max}$ ± SD (h)** | **$T_{1/2}$ ± SD (h)** | **$AUC_{0\text{-}last}$ ± SD (ng*h/mL)** | **$AUC_{0\text{-}inf}$ ± SD (ng*h/mL)** |
| **707GO** | **0.05 mg/kg (n=3)** | 159 ± 33 | 0.67 ± 0.29 | 1.54 ± 0.08 | 466 ± 90 | 481 ± 92 |
| | **0.1 mg/kg (n=3)** | 365 ± 83 | 1.00 ± 0.00 | 1.72 ± 0.57 | 1211 ± 559 | 1299 ± 678 |
| | **0.25 mg/kg (n=3)** | 674 ± 155 | 1.00 ± 0.00 | 1.66 ± 0.13 | 2375 ± 700 | 2486 ± 754 |
| | **1 mg/kg (DIO) (n=2)** | 4590 | 1.00 | 2.59 | 13973 | 13985 |
| **710GO** | **1 mg/kg (DIO) (n=2)** | 3935 | 0.75 | 2.84 | 17776 | 17853 |
| **EBMC-03-47** | **1 mg/kg (n=3)** | 1161 ± 221 | 1.33 ± 0.58 | 2.04 ± 0.67 | 4418 ± 27 | 4453 ± 30 |
| | **3 mg/kg (n=3)** | 3810 ± 118 | 1.33 ± 0.58 | 2.48 ± 0.32 | 14213 ± 985 | 14225 ± 985 |
| | **6 mg/kg (n=3)** | 12437 ± 1983 | 1.17 ± 0.76 | 2.45 ± 0.13 | 49831 ± 5233 | 49873 ± 5244 |
| **EBMC-40-187** | **4 mg/kg (n=3)** | 4862 ± 291 | 0.83 ± 0.29 | 2.49 ± 0.02 | 19885 ± 1459 | 19907 ± 1465 |
| | **8 mg/kg (n=3)** | 11395 ± 3441 | 1.33 ± 0.58 | 2.23 ± 0.06 | 43764 ± 12697 | 43794 ± 12708 |
| | **15 mg/kg (n=3)** | 23046 ± 3450 | 1.00 ± 0.00 | 2.48 ± 0.06 | 85274 ± 7941 | 85351 ± 7958 |
| **Setmelanotide** | **0.5 mg/kg (n=3)** | 767 ± 104 | 0.83 ± 0.29 | 4.01 ± 0.1 | 3105 ± 132 | 3146 ± 136 |
| | **1 mg/kg (n=3)** | 1472 ± 69 | 0.67 ± 0.29 | 3.89 ± 0.16 | 6266 ± 222 | 6340 ± 225 |
| | **3 mg/kg (n=3)** | 4642 ± 1071 | 0.67 ± 0.29 | 3.65 ± 0.24 | 19971 ± 5145 | 20164 ± 5202 |

**Supplementary Table 3 |** Pharmacokinetic parameters in plasma of melanocortin agonist peptides after oral gavage in NHPs at various dose concentrations. All data are presented as mean ± SD.

| | | PK Parameters in Cynomolgus Macaques | | | | |
|---|---|---|---|---|---|---|
| **Compound** | **Oral Gavage Dose** | **$C_{max}$ ± SD (ng/mL)** | **$T_{max}$ ± SD (h)** | **$T_{1/2}$ ± SD (h)** | **$AUC_{0\text{-}last}$ ± SD (ng*h/mL)** | **$AUC_{0\text{-}inf}$ ± SD (ng*h/mL)** |
| **707GO** | **10 mg/kg (n=3)** | 129 ± 25 | 1.67 ± 0.58 | 1.75 ± 0.31 | 387 ± 13 | 410 ± 1 |
| | **10 mg/kg (DIO) (n=4)** | 374 ± 365 | 1.88 ± 1.55 | 7.06 ± 3.08 | 1512 ± 847 | 1662 ± 843 |
| | **30 mg/kg (n=3)** | 193 ± 11 | 1.5 ± 0.87 | 6.44 ± 0.87 | 798 ± 209 | 831 ± 215 |
| **710GO** | **3 mg/kg (n=3)** | 66 ± 51 | 1.67 ± 0.58 | 1.5 ± 0.23 | 207 ± 162 | 216 ± 172 |
| | **10 mg/kg (n=3)** | 133 ± 39 | 1.33 ± 0.58 | 1.84 ± 0.43 | 503 ± 167 | 552 ± 211 |
| | **10 mg/kg (DIO) (n=4)** | 101 ± 88 | 1.5 ± 0.58 | 5.64 ± 2.66 | 541 ± 440 | 581 ± 460 |
| | **30 mg/kg (n=3)** | 248 ± 126 | 2.00 ± 0.00 | 2.47 ± 1.53 | 1006 ± 456 | 1041 ± 437 |
| | **60 mg/kg (n=3)** | 399 ± 164 | 2.33 ± 1.53 | 4.02 ± 0.92 | 1985 ± 856 | 2014 ± 857 |

**Supplementary Table 4 |** Calculated oral bioavailability of 707GO and 710GO in NHPs.

| | PO Dose (mg/kg) | Oral Bioavailability (%F) |
|---|---|---|
| 710GO | 10 | 0.75% |
| | 30 | 0.47% |
| 707GO | 10 | 0.39% |
| | 30 | 0.26% |

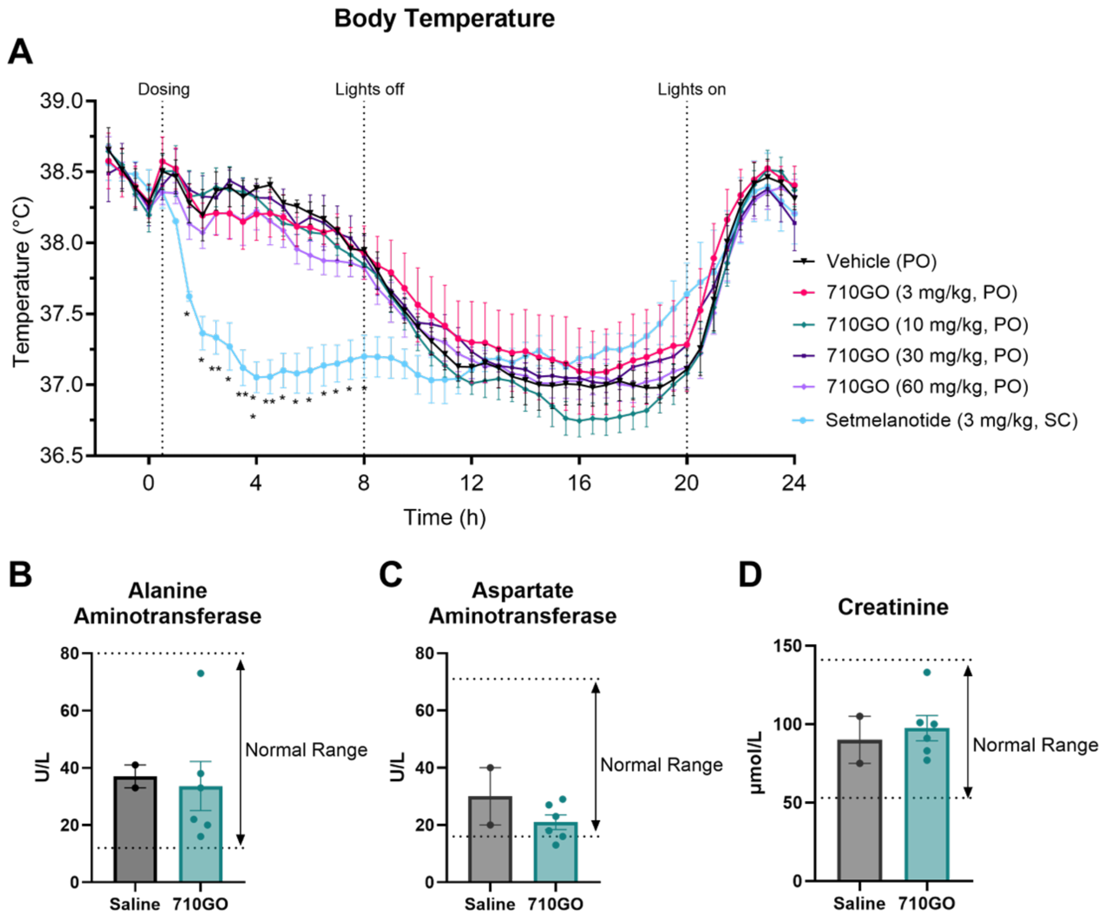


**Supplementary Fig. 11 |** Effects of 710GO on body temperature, alanine aminotransferase, aspartate aminotransferase, and creatinine levels in DIO NHPs. **A**) Body temperature in response to oral administration of 710GO (n=5, per dose group) at doses of 3, 10, 30, and 60 mg/kg and subcutaneous administration of setmelanotide (light blue circle, n=3) at a dose of 3 mg/kg over a 24-hour monitoring period. **B**) Alanine aminotransferase levels after 15 weeks of orally administered 710GO (green circle, n=5) at a dose of 10 mg/kg compared to vehicle (saline, black circle, n=2). **C**) Aspartate aminotransferase levels after 15 weeks of oral 710GO (green circle, n=5) administration at a dose of 10 mg/kg compared to vehicle (saline, black circle, n=2). **D**) Creatinine levels after 15 weeks of oral 710GO (green circle, n=5) administration at a dose of 10 mg/kg compared to vehicle (saline, black circle, n=2). Statistical significance in (A) vs vehicle using Two-Way ANOVA, both followed by Dunnett's multiple comparison test (*p < 0.05, **p<0.01).

A

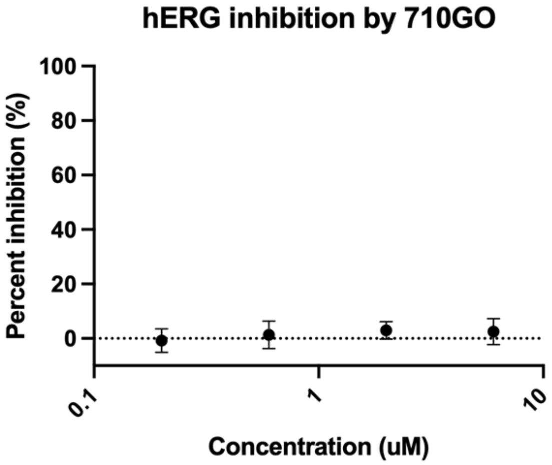


B

| $CL_{int}$ (mic) (µL/min/mg) | $CL_{int}$ (liver) (mL/min/kg) |
|---|---|
| < 9.6 | < 8.6 |

C

| Species | NOAEL (mg/kg) |
|---|---|
| Dog – Male | 300 |
| Dog – Female | 100 |
| Rat – Male | 400 |
| Rat – Female | 400 |

**Supplementary Fig. 12 |** 710GO exhibits safety in hERG, clearance, and AEL studies. **A**) Inhibition of hERG expressed in CHO cells by 710GO. Percent inhibition is relative to the positive control moxifloxacin. **B**) Microsomal and liver clearance values calculated for 710GO incubated at 10 µM with microsomes and hepatocytes, respectively. **B**) Microsomal and liver clearance values calculated for 710GO incubated at 10 µM with human liver microsomes and human hepatocytes, respectively. **C**) Dose levels with no adverse effects (NOAELs) in male and female rodents and canines from 28-day GLP-tox studies.

#### Supplemental Methods:

##### Radioligand Binding study

For radioligand binding assay, competition binding is performed in duplicate in the wells of a 96 well plate (Master Block, Greiner, 786201) containing binding buffer (optimized for each receptor), membrane extracts (amount of protein/well optimized for each receptor), radiotracer (final concentration optimized for each receptor) and test compound. Nonspecific binding is determined by co-incubation with 200-fold excess of cold competitor. The samples are incubated in a final volume of 0.1 mL at a temperature and for a duration optimized for each receptor and then filtered over filter plates. Filters are washed six times with 0.5 mL of ice-cold washing buffer (optimized for each receptor) and 50 µL of Microscint 20 (Packard) are added in each well. The plates are incubated 15 min on an orbital shaker and then counted with a TopCount™ for 1 min/well.

##### NHP acute feeding study

Acute feeding studies in lean male cynomolgus monkeys were conducted at WuXi AppTec. Animals in the fasted groups were fasted overnight. Vehicle (300 mM mannitol in sWFI), EBMC-40-187 (15 mg/kg), EBMC-03-47 (6 mg/kg), and combination EBMC-40-187 + EBMC-03-47 (15 mg/kg + 6 mg/kg) were each administered to Cynomolgus monkeys (n=3, per dose group) via SC injection per the study design. Food consumption was measured at 0.5, 1, 1.5, 2, 3, 4, 24 h post-first dosing (Day 1).

##### Rat husbandry

Male Sprague-Dawley (SD) rats were supplied by Beijing Vital River Laboratory Animal Technology Co., Ltd. The animals were confirmed to be healthy before being assigned to the study. The room(s) was controlled and monitored for relative humidity (targeted mean range 40% to 70%) and temperature (targeted mean range 20°C to 26°C) with 15 or above air changes/hour. The room was on a 12-hour light/dark cycle except when interruptions were necessitated by study activities. Fresh drinking water (reverses osmosis) was available to all animals, *ad libitum*. Certified animal diet was available to all animals when not fasting, *ad libitum*. For fasted animals, animals were fasted at least 12 hours prior to the administration. Certified rodent diet food was withheld until 4 hours post-dose. The fasting time did not exceed 20 hours.

**SD-DIO rat husbandry**

SD-DIO rats are purchased from Beijing Vital River. From 1 week to 6 weeks of age, the animals were given a chow diet, and then they were given a 60% high-fat diet. After arriving at the facility, the animals continued the 60% high-fat diet. The animals are raised in the animal feeding room under strict environmental controls, with the temperature maintained at 20-24°C and humidity maintained at 40-70%. Temperature and relative humidity will be monitored daily. An electronic time-controlled lighting system was used to provide a 12-hour light/12-hour dark cycle (lights off from 19:00 to 7:00) Rats will be housed individually per plastic cage, which is in accordance with the National Research Council “Guide for the Care and Use of Laboratory Animals”. Enrichment toys will be provided. Rats will be fed a high fat diet and fresh water during acclimation; animals will be acclimated in the testing facility for three days.

**SD-DIO Rat acute feeding study**

Acute rat feeding studies were performed at WuXi AppTec. SD-DIO rats were administered vehicle (300 mM mannitol in sWFI) SC injection from Day -5 to Day -1. On day 0, rats were randomized into groups based on body weight and food intake data. On day 0, rats (n=6, per dose group) received a single SC dose of either vehicle or test compound(s) according to the study design: Low dose MC3R (EBMC-40-187, 10 mg/kg), medium dose MC3R (EBMC-40-187, 30 mg/kg), high dose MC3R (EBMC-40-187, 60 mg/kg) or MC4R (EBMC-03-47, 10 mg/kg). On day 0 to day 1, post-dosing food intake was recorded at 1, 4, 8, and 24 hours following compound administration. Body weight and food intake were recorded daily.

**SD-DIO Rat sub-chronic feeding study**

Sub-chronic rat feeding studies were performed at WuXi AppTec. SD-DIO rats (n=6, per dose group) were administered vehicle (300 mM mannitol) or test compounds as a once daily SC injection according to the following study design. Single test article groups: *Days 1-7*: low dose MC3R (EBMC-40-187, 5 mg/kg), high dose MC3R (EBMC-40-187, 15 mg/kg), MC4R (EBMC-03-47, 10 mg/kg), *Days 8-14*: low dose MC3R (EBMC-40-187, 10 mg/kg), high dose MC3R (EBMC-40-187, 30 mg/kg), MC4R (EBMC-03-47, 10 mg/kg), Combination group: *Day 7-8:* MC3R + MC4R (EBMC-40-187, 30 mg/kg + EBMC-03-47, 10 mg/kg). *Day 9-14:* MC3R + MC4R (EBMC-40-187, 10 mg/kg + EBMC-03-47, 10 mg/kg). Combination group Day 7 was aligned with Day 0 of the other dose groups. Cumulative food intake and body weight were measured for five days prior to compound administration and daily post-dosing.

**SD-DIO Rat acute feeding study (dual agonism)**

Rat feeding studies were performed at WuXi AppTec. Male SD-DIO rats were administered vehicle (300 mM mannitol in sWFI) or test compounds as once daily SC injection according to the following study design: 707GO (3 mg/kg), 710GO (3 mg/kg).

Cumulative food consumption was measured 1, 4, 8, 24 hours post dose and then daily for 17 days. Caloric consumption and body weight were measured daily.

**SD-DIO Rat sub-chronic feeding study (dual agonism)**

Rat feeding studies were performed at WuXi AppTec. Male SD-DIO rats were fed a 45% high fat diet prior to study initiation and maintained on the high fat diet for the duration of the study. SD-DIO rats were administered vehicle (saline) by daily SC injection from Day -4 to Day -1. Rats were administered vehicle (saline) or test compounds as once daily SC injection according to the following study design (n=4, vehicle; n=5, per dose group); Days 1-7: 707GO (0.5 mg/kg), 710GO (0.5 mg/kg), Days 8-17: 707GO (1.0 mg/kg), 710GO (1.0 mg/kg). Cumulative food consumption was measured 1, 4, 8, 24 hours post dose and then daily for 17 days. Body weight was measured daily.

**Pharmacokinetic studies in rats**

Pharmacokinetic studies in male Sprague-Dawley Rats were conducted at WuXi AppTec. Rats (n=3, per dose group) were administered test compounds by subcutaneous injection or by oral gavage. The vehicle was physiological saline. For all groups, animals were fasted overnight through 4 hours post dosing. Plasma and CSF samples were collected at 0.083, 0.25, 0.5, 1, 2, 4, 6, 8, 10 and 24 hours post-dose. *Blood sample collection:* Blood (at least 0.1 mL) was collected at each time point via jugular vein puncture from each study animal. All blood samples were transferred into commercial tubes containing K2-EDTA. After blood was collected, the samples were placed on wet ice and processed for plasma within 60 min of collection by centrifugation at approximately 4°C, 3200×g for 10 minutes, and then quickly frozen over dry ice and kept at -60°C or lower until LC-MS/MS analysis. Concentrations of test compounds in samples were determined by a liquid chromatography tandem mass spectrometry (LC-MS/MS) method. *CSF sample collection:* CSF samples were collected at each time point from cisterna magna while animal was under anesthesia with isoflurane. At the sampling time point, needle was inserted at base of skull and at least 20 µL CSF was dropped into EP tube from each rat. The volume of CSF was recorded, and the collected CSF should be quick-frozen over dry ice and stored at -60°C or lower until LC-MS/MS analysis. The linear/log trapezoidal rule was applied in obtaining the PK parameters. Individual plasma and CSF concentration values that were below the lower limit of quantitation (LLOQ) were excluded from the PK parameter calculation. The nominal dose levels and nominal sampling times were used in the calculation of all pharmacokinetic parameters.

**Pharmacokinetic studies in Cynomolgus macaques**

Pharmacokinetic studies in non-naïve male cynomolgus monkeys were conducted at WuXI AppTec. Animals were administered test compounds by subcutaneous injection or oral gavage (n=3, per dose group). The vehicle was physiological saline. For all groups, animals were fasted overnight through 4 hours post dosing. Plasma samples were collected at 0.083, 0.25, 0.5, 1, 2, 4, 6, 8 and 24 hours post dose. Concentrations of test compounds in plasma samples were determined by a liquid chromatography tandem mass spectrometry (LC-MS/MS) method. At least 0.5 mL blood was collected at each time point via peripheral vessel from each study animal. All blood samples were transferred into low binding EP tubes (0.5 M K2 EDTA was pre-added as a ratio of 50:1 for blood: anticoagulant) and placed on wet ice. Plasma samples were then prepared by centrifuging the blood samples at approximately 2°C to 8°C, 3200×g for 10 min within one hour of collection. The plasma samples about 0.2 mL were divided to approximate 0.1 mL×2 aliquots, transferred into labeled polypropylene micro-centrifuge tubes and stored frozen in a freezer set to maintain -60°C or lower until bio-analysis. The plasma concentrations of test compounds in study animals were subjected to a non-compartmental pharmacokinetic analysis by using the Phoenix WinNonlin software (version 8.3.5, Certara). The linear/log trapezoidal rule was applied in obtaining the PK parameters. Individual plasma concentration values that were below the lower limit of quantitation (LLOQ) were excluded from the PK parameter calculation. The nominal dose levels and nominal sampling times were used in the calculation of all pharmacokinetic parameters.

**Oral bioavailability studies in Cynomolgus macaques**

Pharmacokinetic studies in non-naïve male cynomolgus monkeys were conducted at WuXi AppTec. Animals were administered test compounds by intravenous bolus or oral gavage (n=3, per dose group). Test compounds were formulated in water for injection (sWFI). Plasma samples were collected at 0.083, 0.25, 0.5, 1, 2, 4, 6, 8 and 24 hours post dose. Concentrations of test compounds in plasma samples were determined by a liquid chromatography tandem mass spectrometry (LC-MS/MS) method. At least 0.5 mL blood was collected at each time point via peripheral vessel from each study animal. All blood samples were transferred into low binding EP tubes (0.5 M K2 EDTA was pre-added as a ratio of 50:1 for blood: anticoagulant), and placed on wet ice. Plasma samples were then prepared by centrifuging the blood samples at approximately 2°C to 8°C, 3200×g for 10 min within one hour of collection. The plasma samples about 0.2 mL was divided to approximate 0.1 mL×2 aliquots, transferred into labeled polypropylene micro-centrifuge tubes and stored frozen in a freezer set to maintain -60°C or lower until bio-analysis. The plasma concentrations of test compounds in study animals were subjected to a non-compartmental pharmacokinetic analysis by using the Phoenix WinNonlin software (version 8.3.5, Certara). The linear/log trapezoidal rule was applied in obtaining the PK parameters. Individual plasma concentration values that were below the lower limit of quantitation (LLOQ) were excluded from the PK parameter calculation. The nominal dose levels and nominal sampling times were used in the calculation of all pharmacokinetic parameters. Oral bioavailability (%F) was calculated using $AUC_{0\text{-}last}$ (ng.h/mL) and nominal dose in the following formula.

$$\%\mathbf{F} = \mathbf{100} \times \frac{\boldsymbol{AUC_{po}} \times \boldsymbol{dose_{iv}}}{\boldsymbol{AUC_{iv}} \times \boldsymbol{dose_{po}}}$$

**hERG study**

CHO-hERG cells were obtained from Sophion Biosciences (Ballerup, Denmark), subcultured and frozen in WuXi AppTec (Suzhou) Co., Ltd. Complete medium was F12 medium, supplemented with 10% fetal bovine serum, 1% Geneticin® selective antibiotic (G418), and 89 µg/mL Hygromycin B (HB). Recovery medium was F12 medium with 10% fetal bovine serum. CHO-hERG cells were cultured in a humidified incubator of 5% CO2 (4% to 8%) in air at 37°C (±2°C). Exponentially growing CHO-hERG cells were collected and suspended in extracellular solution for use. The hERG current was recorded at 35.0°C to 37.0°C using whole-cell patch-clamp techniques. Output signals from the patch-clamp amplifier were digitized and low-pass filtered at 2.9 KHz. The recording was controlled with Patchmaster Pro software. A micropipette filled with intracellular solution (ICS, composition: KOH, 31.25 mM; KCl, 120 mM; MgCl2, 1.75 mM; CaCl2, 5.374 mM; EGTA, 10 mM; Na2ATP, 4 mM and HEPES, 10 mM) was used as recording electrode in the manual patch-clamp study. The voltage protocol was 3900 ms in duration, and was repeated every 5 s. The voltage “ramp down” phase was 100 ms in duration, from +40 mV to -80 mV (hence a voltage change of 1.2 V/s). The peak current amplitude during the “ramp down” phase was recorded. Peaks were monitored for at least 25 sweeps until stable. Each recording was completed with the application of a supramaximal concentration of the selective blocker (E-4031, 1 μM), which allow to assess the contribution of background currents, and the perfusion duration was at least 2 minutes. In the definitive hERG assay, four replicate cells at four concentrations each of test article working solutions of 0.2, 0.6, 2 and 6 μM were tested. The average peak current amplitude of the last ten sweeps of every 4 minutes was used to evaluate the current run-down or run-up. All the values were normalized to that of the first 4 minutes and presented as percentage. Less than 15% current run-down or run-up during the measurement period was considered acceptable. Moxifloxacin was used as the positive control article to evaluate the reliability of the test system. When acquiring the whole-cell configuration, a holding potential (e.g., -80 mV) was applied while membrane parameters were collected ($C_m$, $R_m$ and $R_s$). Input resistance ($R_{input}$) was monitored and calculated as follows: ($R_{input}$) = (V-90mV- V-80mV)/ (I-90mV- I-80mV).

**Microsome and Liver Intrinsic Clearance Study**

Metabolic stability, expressed as percent of the parent compound remaining, was calculated by comparing the peak area of the compound at the time point relative to that at time-0. Target compound was incubated at 0, 15, 30, 45, and 60 minutes at 37°C, and detected by HPLC-MS/MS. The half-life ($T_{1/2}$) was estimated from the slope of the initial linear range of the logarithmic curve of compound remaining (%) vs. time, assuming the first-order kinetics. The apparent intrinsic clearance ($CL_{int}$, in μL/min/mg or mL/min/kg) was calculated according to the following formula: CLint= 0.693 $T_{1/2}$*(mg protein/μL or million cells/μL).
